# p53 preserves genome stability in dividing cells under hypo-osmotic stress

**DOI:** 10.64898/2025.12.17.694541

**Authors:** Bin Sheng Wong, Alokendra Kumar Ghosh, Hongyue Cui, Keming Li, Blake A. Johnson, Kriti Sethi, Yaelim Lee, Jin Zhu, Sean X. Sun, Rong Li

## Abstract

Cells respond to frequent osmotic stress to maintain structural integrity and function. Whereas acute volume regulation is well-known, the cellular response to prolonged osmotic stress and its long-term consequences remain poorly understood. In this study, we investigate how lasting hypo-osmotic stress influences cell growth and division and the associated cellular response. We observe that cells undergoing their first division under hypo-osmotic stress experience delayed mitotic entry, lengthened mitotic duration, elevated mitotic errors and aneuploidy formation. However, continued cell division is halted through a p53/p21-dependent G_1_ arrest. p53 activation occurs post-mitotically and is associated with hyper-condensed chromatin and impaired nuclear expansion in daughter cells after mitotic exit. We show that the altered nuclear size dynamics lead to nuclear accumulation of p53 and activation of downstream gene expression. These findings uncover a mechanosensitive, p53-mediated mechanism that safeguards genome integrity by suppressing erroneous mitosis under hypo-osmotic stress.

## Introduction

Cell volume and osmotic homeostasis are tightly regulated processes critical for cellular architecture and function. Whereas steady-state cell volume correlates with biomass and cell cycle stage^1^, acute changes in cell volume are primarily governed by osmotic gradients across the semi-permeable plasma membrane^2^. When these gradients are disrupted due to altered extracellular or intracellular solute concentrations, water rapidly enters or leaves the cell, leading to abrupt changes in cytoplasmic water content, protein concentration and cell volume^3^. To survive, cells must respond and correct such deviations within minutes by activating specific membrane-bound channels and pumps to modulate osmolyte and water flux^4,5^. Nevertheless, despite swift volume recovery, the intracellular ionic and osmotic environment inevitably changes under sustained osmotic stress, which could lead to profound long-term consequences for cell physiology.

Systemic osmolality of many organisms is maintained within a narrow range of approximately 300 mosmol/kg by the renal and endocrine systems^6^. However, cells within certain tissues, such as kidney, skin, lungs, eyes and intestines, frequently experience osmotic fluctuations in their microenvironment that can last from hours to days, which can cause potential tissue damage and diseases^7–10^. Hyper-osmotic stress, where the external environmental solute concentration is higher than the cell interior, is well known to be linked to diabetes, cystic fibrosis, burns, kidney and neurodegenerative diseases^11^. The opposite hypo-osmotic stress is likewise prevalent, with one of the major causes being hyponatremia, where abnormally low blood sodium level causes water influx and cellular swelling^12^. Hypo-osmotic stress is also associated with pathological conditions such as cerebral edema, tissue injury, and inflammation^13–16^. Accumulation of inflammatory mediators during chronic inflammation can influence ion and fluid transport and increase vascular permeability, leading to excess edematous fluid that can create localized regions of hypotonicity in inflamed tissues^17,18^. Moreover, hypo-osmotic stress has also been shown to promote cytokine production^19,20^, suggesting a self-reinforcing cycle between chronic inflammation and hypo-osmotic stress.

Aside from osmolarity perturbation in the tissue microenvironment, hypo-osmotic stress can also arise from changes in intracellular properties and ionic concentrations. This is particularly evident in hydropic swelling, where cells swell from excessive intracellular ion and water accumulation, leading to cytoplasmic vacuolation despite iso-osmotic extracellular conditions^21^. This phenomenon is typically caused by cell membrane damage and ion pump failure during tissue injury in conditions such as liver hepatitis, acute tubular necrosis and myocardial infarction^22–24^. Beyond injury, our work revealed that aneuploid cells experience hypo-osmotic-like stress due to increased intracellular osmolarity driven by proteome imbalance^25,26^. As chronic inflammation predisposes tissues to neoplasia and aneuploidy is a common genomic alteration characteristic of most cancers^27,28^, we hypothesize that a vicious cycle between osmotic stress and chromosome instability could be an underlying driver of inflammation-associated cancers.

Testing the above hypothesis requires a thorough understanding of the cellular response and adaptive process to osmotic stress at physiologically relevant time scales. In this study, we examined how cells respond to chronic hypo-osmotic stress and the consequences on cell proliferation and genome stability. Our results show that prolonged hypo-osmotic stress disrupts mitotic fidelity. Interestingly, non-transformed cells subsequently stop proliferation and suppress error propagation through p53 activation triggered by impaired post-mitotic nuclear expansion. Our study thus uncovers a critical mechanosensitive safeguard that protects cells from osmotic stress-induced genome instability.

## Results

### Hypo-osmotic stress delays mitotic progression and impairs chromosome segregation

We first used live imaging to follow cell division cycle under sustained hypo-osmotic stress. Hypo-osmotic cell culture media were formulated by diluting standard culture medium with deionized water. Corresponding isotonic controls, whereby the same culture medium was diluted with phosphate-buffered saline (PBS) at the same proportions, were used to account for nutrient dilution. In most experiments, a 1:1 ratio of medium to water (Hypo) at 136.0 ± 5.6 mmol/kg was used and compared to a 1:1 ratio of medium to PBS iso-osmotic control (Iso) at 294.7 ± 2.1 mmol/kg and an undiluted medium control (Medium) at 309 ± 4.6 mmol/kg. RPE-1, an hTERT-immortalized, chromosomally stable, near-diploid cell line, served as the primary cell model in this study, although key findings were also validated in other cell models as described below.

RPE-1 cells expressing H2B-mNeonGreen were treated with Medium, Iso, or Hypo and observed for 24 hr (Movie 1). Although most cells were able to enter and complete mitosis with seemingly normal chromosome segregation, approximately 30% of mitosis under Hypo exhibited signs of abnormalities, significantly higher than the basal error rate of 5% in Medium or Iso (Figure 1A-B). These abnormalities included lagging chromosomes (LC), anaphase bridges (AB), unaligned chromosomes, multipolar mitosis and mitotic slippage (Supplementary Figure 1A). In many instances, mitotic errors, typically LC, led to the formation of micronuclei in daughter cells (Figure 1A). Most erroneous mitoses involved a single type of error, with LC and AB occurring at comparable frequencies (Figure 1C).

**Figure 1:**
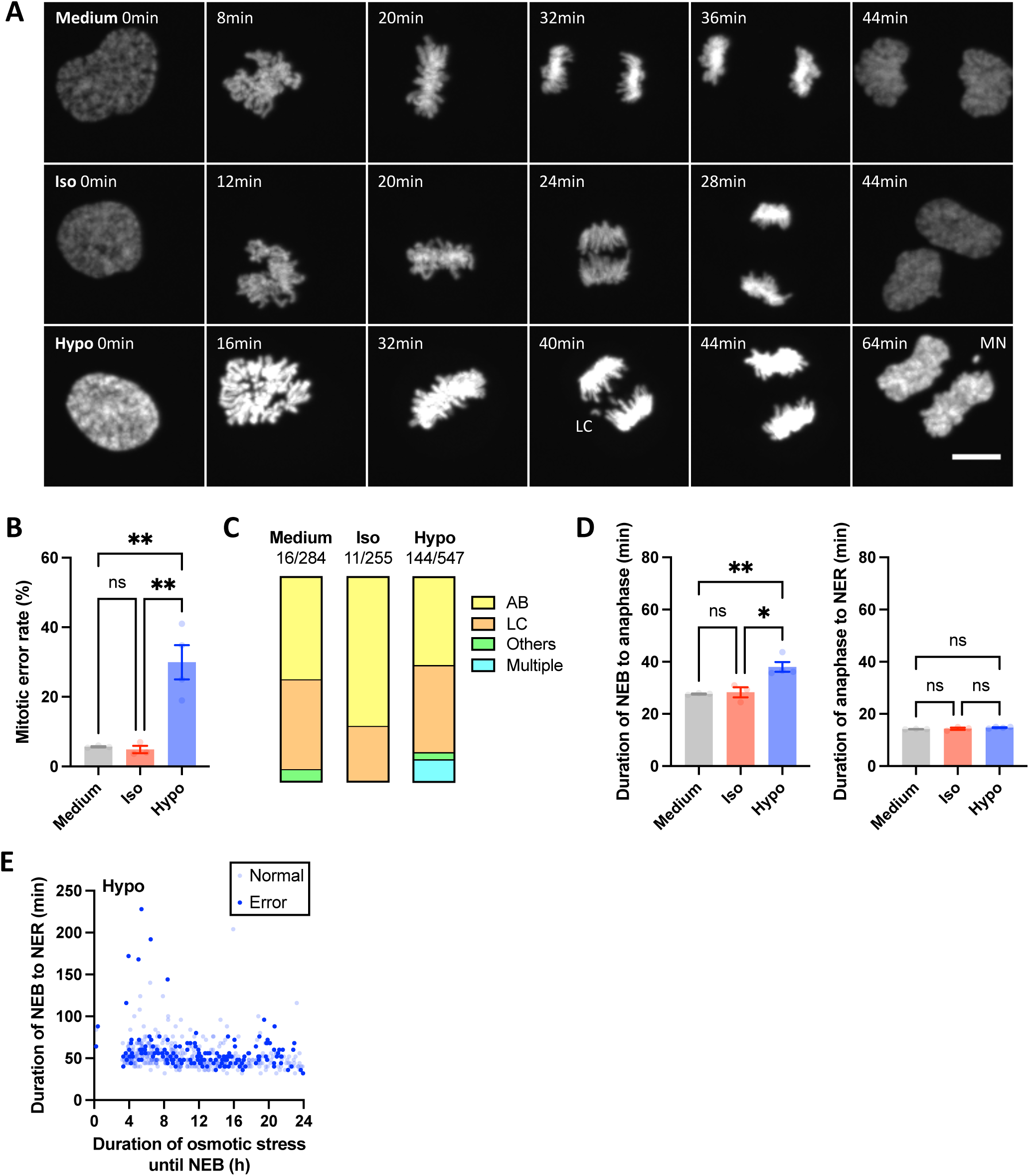
Hypo-osmotic stress delays mitotic progression and impairs chromosome segregation. **(A)** Time-lapse montages of RPE-1 H2B-mNeonGreen cells undergoing mitosis under control (Medium), iso-osmotic (Iso, 1:1 medium:PBS), or hypo-osmotic (Hypo, 1:1 medium:water) conditions. Images were aligned to nuclear envelope breakdown (NEB, t = 0 min) and depict key stages of mitosis: prophase, metaphase, anaphase, telophase and nuclear envelope reformation (NER). Mitotic abnormalities such as lagging chromosomes (LC) and micronuclei (MN) are indicated. Scale bar = 5 µm. **(B)** Frequency of mitotic errors under different osmotic conditions. **(C)** Classification of mitotic errors observed under different osmotic conditions: anaphase bridges (AB), LC, others (unaligned chromosomes, multipolar mitosis or mitotic slippage), and multiple concurrent errors. The total number of erroneous over all mitoses observed within 24 hr of osmotic treatment across all experimental replicates (n ≥ 3) is indicated. **(D)** Quantifications of the duration of NEB to anaphase (left), and anaphase to NER (right) under different osmotic conditions. **(E)** Correlation between the duration of NEB to NER (i.e. mitotic duration) and the duration of osmotic stress exposure until NEB for all mitotic events under Hypo condition across all experimental replicates (n≥3). Normal (light) and erroneous (dark) mitoses are differentially labeled. Bar graphs in (B) and (D) represent mean ± SEM from n ≥ 3 independent experiments. For (D), data of multiple cells from each independent experiment were averaged to generate a single biological replicate value. Statistical significance was assessed via one-way ANOVA for (B) and (D). ns represents not significant, * represents p ≤ 0.05, ** represents p ≤ 0.01, *** represents p ≤ 0.001 and **** represents p ≤ 0.0001.

The duration of mitosis was significantly lengthened under Hypo (Figure 1D and Supplementary Figure 1B). Specifically, the time from nuclear envelope breakdown (NEB) to anaphase onset was prolonged in Hypo, but not the time from anaphase to nuclear envelope reformation (NER). Mitotic duration (i.e. NEB to NER) was not correlated with the duration of osmotic stress until mitotic initiation at NEB, implying that the extent of mitotic prolongation was not influenced by the duration of stress exposure (Figure 1E and Supplementary Figure 1C). Both normal and erroneous mitoses shared similar mitotic durations and occurred throughout the stress period. Interestingly, there was a prominent delay of around 4 hr between the start of Hypo treatment and the bulk of first mitotic entries, which was absent in Medium and Iso, suggesting that mitotic entry was paused momentarily and delayed after Hypo exposure (Figure 1E and Supplementary Figure 1C). Taken together, these findings reveal that hypo-osmotic stress delays cell cycle progression and compromises mitotic fidelity.

### Hypo-osmotic stress triggers G_1_ cell cycle arrest to limit cell proliferation

To evaluate the consequences of defective mitosis under hypo-osmotic stress, we followed the proliferation and cell cycle progression of osmotically treated cells for 72 hr. As a whole population, Hypo cells exhibited significantly slower growth compared to both Medium and Iso cells despite the same initial seeding density (Figure 2A and Supplementary Figure 2A). The growth of Iso cells was slightly diminished relative to Medium cells, likely due to nutrient dilution. The decreased proliferation of Hypo cells, compared to that of Iso cells, was further evidenced by significant reductions in EdU-positive (Figure 2B) and Ki67-positive (Supplementary Figure 2B) cell populations. Flow cytometry-based cell cycle profiling further suggested that Hypo cells ceased proliferation (Figure 2C), as demonstrated by the near-complete disappearance of S/G_2_ cell population after 72 hr.

**Figure 2:**
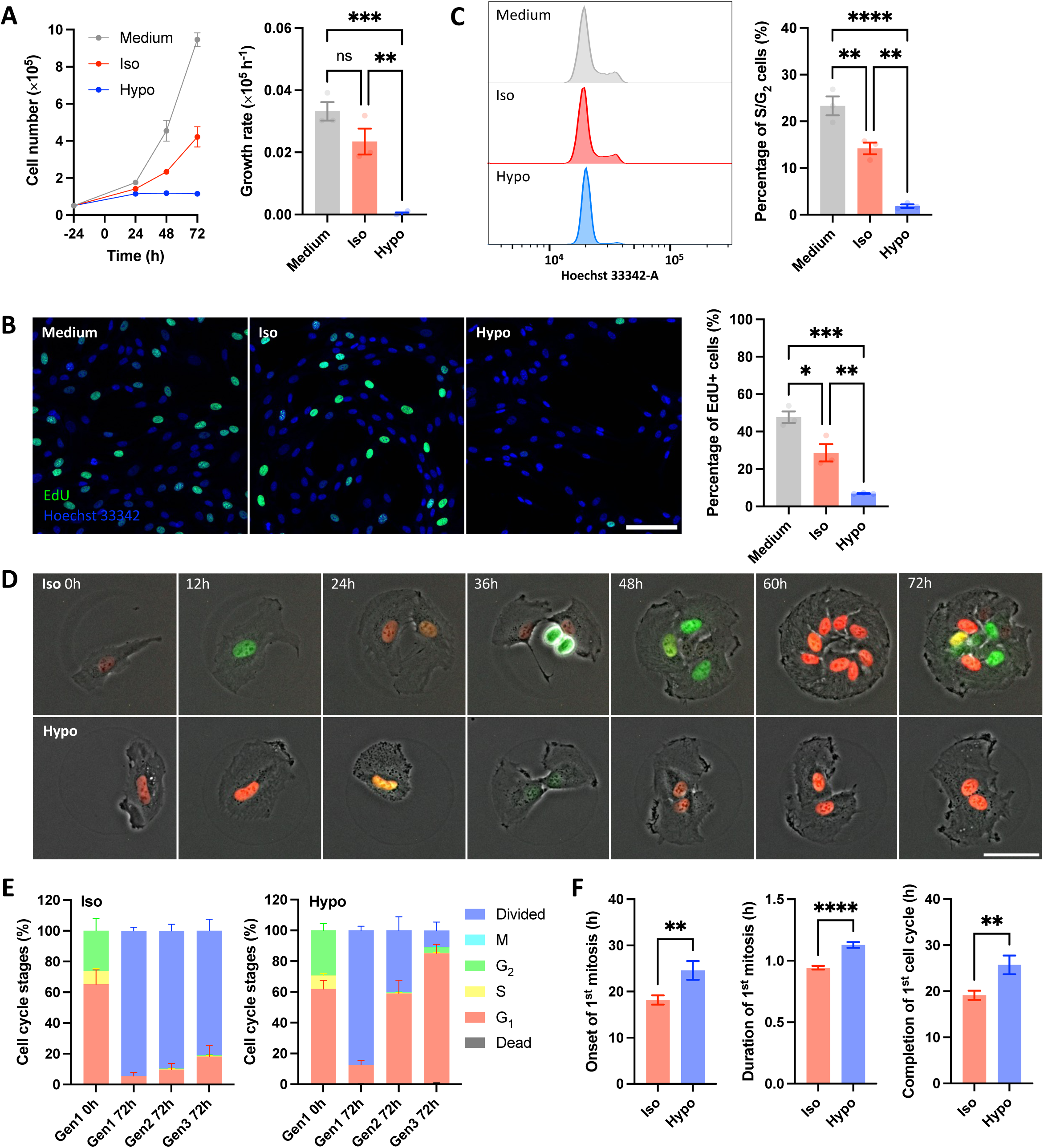
Hypo-osmotic stress triggers G_1_ cell cycle arrest to limit cell proliferation. **(A)** Growth curves of RPE-1 cells under different osmotic conditions (left). The dots and error bars represent mean ± SEM from n ≥ 3 independent experiments. Growth rates were calculated by exponential curve fitting using the least-squares method (right). **(B)** EdU staining (green) showing proliferating cells with newly synthesized DNA after 72 hr of osmotic stress, along with Hoechst 33342 nuclear counterstain (blue). Scale bar = 100 µm (left). Quantification of the percentage of EdU+ cells (right). **(C)** Cell cycle profiles by genome content labeling after 72 hr of osmotic stress (left). Quantification of the percentage of S/G_2_ cells (right). **(D)** 72 hr time-lapse montages of FUCCI-labeled RPE-1 cells on micropatterns under Iso (top) or Hypo (bottom) conditions. (red, G_1_; yellow/orange, S; green, G_2_). Scale bar = 50 µm. **(E)** Cell cycle phase distribution of Gen 1 (initially seeded single cell), Gen 2 (daughter cells from Gen 1 mitosis), and Gen 3 cells (daughter cells from Gen 2 mitosis) at 0 hr and 72 hr under Iso (left) or Hypo (right) conditions. Data represents mean ± SEM from n ≥ 3 independent experiments. **(F)** Quantifications of the time to onset of 1^st^ mitosis (left), the duration of 1^st^ mitosis (middle), and the time to completion of 1^st^ cell cycle (right) under Iso or Hypo conditions. Bar graphs in (A), (B), (C) and (F) represent mean ± SEM from n ≥ 3 independent experiments. For (B), data obtained from multiple fields of view from each independent experiment were averaged to generate a single biological replicate value. Statistical significance was assessed via one-way ANOVA for (A), (B) and (C), and t-tests for (F). ns represents not significant, * represents p ≤ 0.05, ** represents p ≤ 0.01, *** represents p ≤ 0.001 and **** represents p ≤ 0.0001.

In addition to RPE-1 cells, a similar Hypo-induced growth inhibition was also observed in other cell models, including the adherent BJ-5ta fibroblasts and MDCK epithelial cells, the non-adherent Nalm6 suspension leukemia cells, as well as primary mouse colon organoids in three-dimensional culture (Supplementary Figure 2C-D). Furthermore, the degree of growth inhibition was dependent on the strength of hypotonicity: the lower the media osmolality, the greater the decrease in growth rate and the percentage of S/G_2_ population (Supplementary Figure 2E). Interestingly, Hypo-induced cell cycle arrest was reversible: cells treated with Hypo for 72 hr re-entered cell cycle upon switching to Iso but remained arrested under continued Hypo (Supplementary Figure 2F). These results suggest that the growth-inhibitory effect of hypo-osmotic stress is generalizable to diverse cell types, dependent on the strength of hypotonicity, and reversible upon stress withdrawal.

### Hypo-osmotic stress-induced cell cycle arrest occurs after the initial round of cell division

To further characterize the effects of hypo-osmotic stress on cell cycle progression, individual RPE-1 cells stably expressing the FUCCI cell cycle marker^29^ were imaged for 72 hr in Iso or Hypo condition. Cells were seeded onto fibronectin-coated circular micropatterns to facilitate tracking of single cells and their progenies across successive cell divisions. Iso cells underwent multiple rounds of division (Movie 2, Figure 2D), whereas most Hypo cells completed only one or two divisions before becoming G_1_-arrested (Movie 3, Figure 2D). To quantify this phenomenon, cell cycle stages were characterized at each cell division generation (Gen). In the beginning of the experiment (0 hr), initially seeded Gen 1 cells were asynchronous and showed comparable cell cycle distributions in both Iso and Hypo conditions (Figure 2E). At the end of the experiment (72 hr), almost all Gen 1 cells divided under both conditions to produce Gen 2 cells. However, Hypo Gen 2 cells became predominantly G_1_-arrested, with only a minority progressing to Gen 3, of which nearly all subsequently became G_1_-arrested. In contrast, Iso cells maintained high proliferative capacity across generations. This data demonstrates that hypo-osmotic stress permits initial mitotic completion but induces potent and progressive post-mitotic G_1_ arrest.

Additionally, the time taken to initiate and complete mitosis were also quantified. As Gen 1 cells were asynchronous, the time to the onset of first mitosis (i.e. NEB) was quantified and appeared significantly prolonged in Hypo (Figure 2F). Hypo cells also took longer to complete mitosis and the first cell cycle (Figure 2F). Similar prolongations were also observed in the second round of cell cycle for dividing Hypo Gen 2 cells (Supplementary Figure 2G). Overall, these results corroborate with the earlier observations made with H2B-mNeonGreen cells, that hypo-osmotic stress delays mitotic initiation and completion.

### Hypo-osmotic stress downregulates cell cycle-related pathways while activating p53

To investigate the molecular pathways underlying the changes in mitotic cell cycle under hypo-osmotic stress, a time-course RNA sequencing (RNA-seq) experiment was performed using RNA samples collected at 6, 12, 24, 48 and 72 hr of culture under Iso or Hypo. Principal component analysis showed that the transcriptomic profiles of Iso and Hypo cells were clearly distinct (Figure 3A). The effects of stress duration were also detectable, especially in Hypo where a clear trajectory with time was evident.

**Figure 3:**
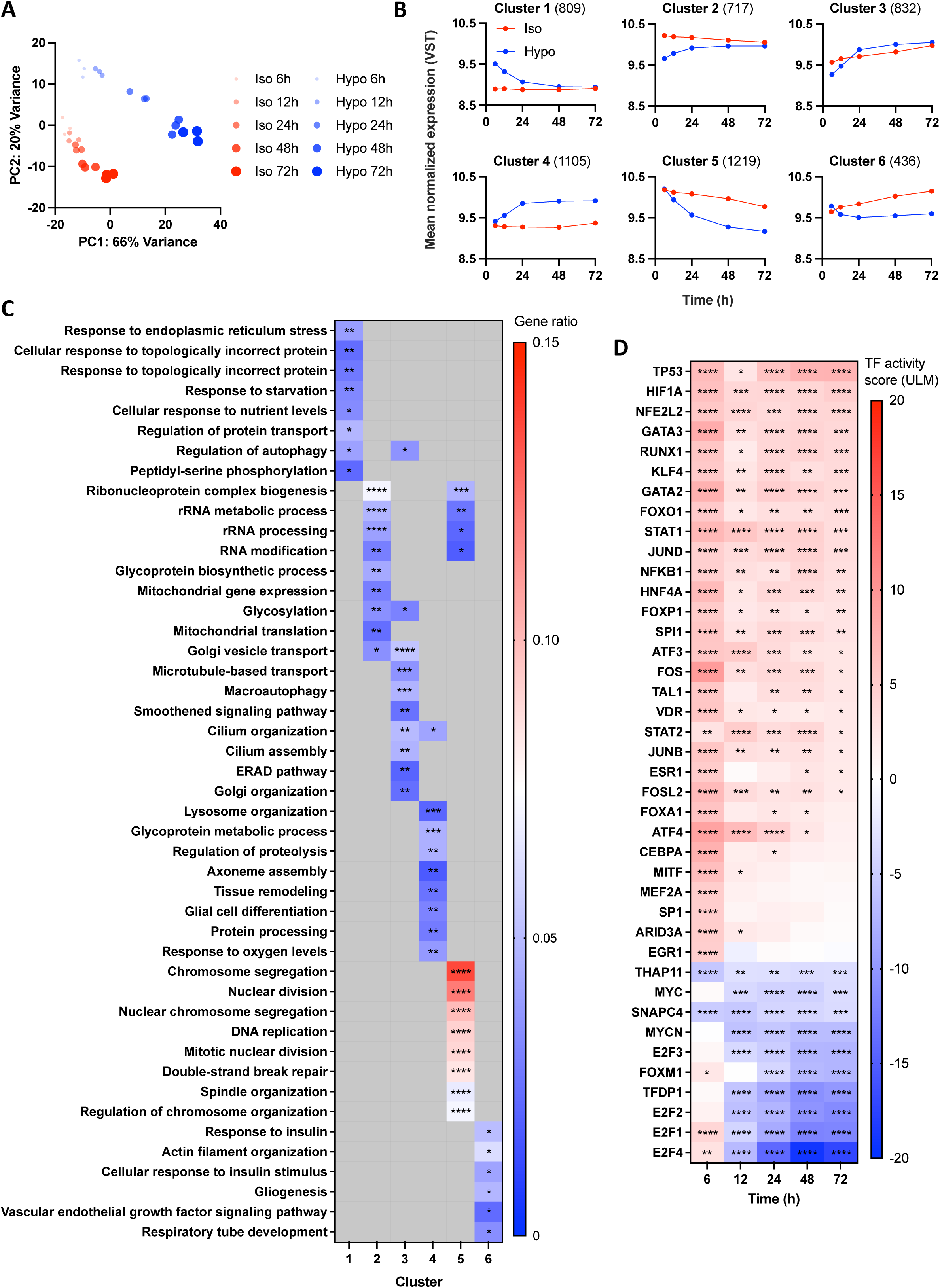
Hypo-osmotic stress downregulates cell cycle-related pathways while activating p53. **(A)** Principal component analysis (PCA) of RNA-seq data from Iso- (red) or Hypo-treated (blue) RPE-1 cells at different timepoints (i.e. 6, 12, 24, 48 and 72 hr). Dot size represents the duration of stress. **(B)** Normalized gene expression profiles of Iso- (red) or Hypo-treated (blue) cells overtime for the six clusters identified by K-means clustering (see Supplementary Figure 3A). The number of genes in each cluster is indicated in parenthesis. Expression values were normalized via Variance Stabilizing Transformation (VST). **(C)** Gene Ontology Biological Process enrichment comparison for the six gene clusters. Top significantly enriched pathways of each cluster are shown. The heatmap represents the gene ratio. **(D)** Transcription factor activity analysis of the time-course RNA-seq data inferred using decoupleR (Univariate Linear Model, ULM). Top 40 transcription factors are shown. The heatmap represents the ULM activity score, with red indicating activation and blue indicating inactivation (Hypo vs Iso contrast). The transcription factors are ordered by decreasing ULM score at 72 hr. Statistical significance for enrichment and activity analyses are represented by adjusted P values, where * represents p ≤ 0.05, ** represents p ≤ 0.01, *** represents p ≤ 0.001 and **** represents p ≤ 0.0001.

We used a likelihood ratio test to identify genes with significant time-dependent changes between conditions using the R package DESeq2^30^. K-means clustering analysis of these genes identified six gene expression clusters with distinct temporal patterns (Figure 3B, Supplementary Figure 3A). Some clusters, such as Cluster 5, exhibited a trend of gradual and progressive gene downregulation over time in Hypo, whereas Cluster 4 displayed a trend of gradual upregulation (Figure 3B). In contrast, some cluster, such as Clusters 1 and 2, captured Hypo-induced transcriptomic changes that were more specific to the earlier time points. Gene Ontology Biological Process enrichment analyses were subsequently conducted on the different clusters (Figure 3C). Notably, cell cycle and mitosis-related terms were prominently represented by Cluster 5 which downregulated progressively with increasing duration of Hypo treatment. Indeed, many genes involved in various stages of cell cycle progression and mitosis were suppressed over time under Hypo, such as cyclin-dependent kinases (*CDK1*), cyclins (*CCNB1*), cell division cycle proteins (*CDC20*), aurora kinases (*AURKA*), condensins (*SMC4*), kinesins (*KIF11*), cytokinesis-associated proteins (*ANLN*) and centromeric proteins (*CENPA*) (Supplementary Figure 3B).

Cell cycle inhibition is most prominently regulated by the tumor suppressor gene p53 in response to a variety of stress conditions^31^. Upon p53 activation, its downstream transcriptional target p21 is induced and serves to inhibit cyclins and CDKs, ultimately inhibiting the E2F family of transcription factors which are master transcriptional regulators of cell cycle progression and DNA synthesis^32^. To investigate the underlying regulatory mechanisms, we inferred transcription factor activities using the decoupleR framework with DoRothEA regulons^33^. This analysis revealed that p53 was activated, while the E2F transcription factors and its dimerization partner DP1 (encoded by *TFDP1*) were inactivated overtime under Hypo treatment (Figure 3D). The involvement of p53 and E2F transcription factors was further implicated by the TRRUST analysis which infers transcriptomic regulatory relationships from differentially expressed genes^34^ (Supplementary Figure 3C).

### p53 activation prevents propagation of hypo-osmotic stress-induced genomic instability

Although p53 mRNA was only marginally elevated after 72 hr of Hypo, the mRNA level of its downstream transcriptional target p21 was markedly increased (Supplementary Figure 4A). Western blot analysis demonstrated that Hypo led to a slight elevation of p53 protein level but a more prominent upregulation of p21 protein (Figure 4A). A similar trend of p53-p21 pathway activation was also observed in mouse colonoids under Hypo (Supplementary Figure 4B).

**Figure 4:**
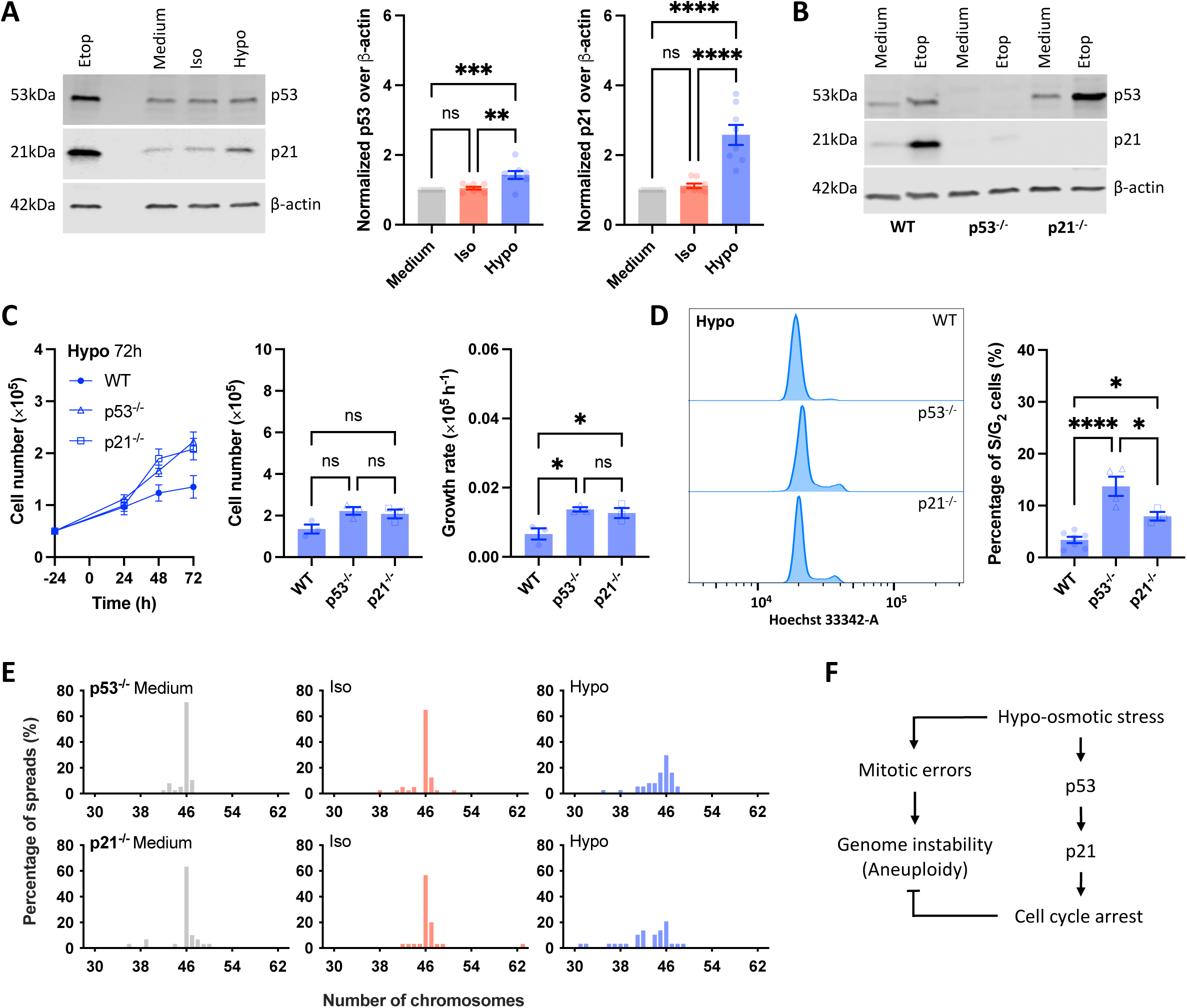
p53 activation prevents propagation of hypo-osmotic stress-induced genomic instability. **(A)** Western blots of p53 and p21 in RPE-1 cells after 72 hr of osmotic stress (left). Treatment with 10 µM Etoposide (Etop) for 24 hr served as a positive control for DNA damage-induced p53 and p21 activation. Quantifications of the protein level of p53 (middle) and p21 (right) normalized to β-actin. **(B)** Knockout validation of p53 (p53^-/-^) and p21 (p21^-/-^) in wild type (WT) RPE-1 cells by CRISPR via western blots. Etop was included as a DNA damage positive control. **(C)** Growth curves of WT, p53^-/-^ and p21^-/-^ cells under Hypo conditions (left). The dots and error bars represent mean ± SEM from n ≥ 3 independent experiments. Quantifications of cell number (middle) and growth rates (right) of WT, p53^-/-^ and p21^-/-^ cells after 72 hr of Hypo treatment. **(D)** Cell cycle profiles by genome content labeling of WT, p53^-/-^ and p21^-/-^ cells after 72 hr of Hypo treatment (left). Quantification of the percentage of S/G_2_ cells (right). **(E)** Chromosome number distribution of p53^-/-^ (top) and p21^-/-^ (bottom) cells after 72 hr of osmotic stress by metaphase spread analysis. Data represents ≥ 30 metaphase spreads from ≥ 3 independent experiments for each experimental condition. **(F)** Model of the proposed role of p53 activation under hypo-osmotic stress. Hypo-osmotic stress induces a p53- and p21-dependent G_1_ cell cycle arrest that halts error-prone mitosis to prevent genome instability. All bar graphs represent mean ± SEM from n ≥ 3 independent experiments. Statistical significance was assessed via one-way ANOVA for (A), (C) and (D). ns represents not significant, * represents p ≤ 0.05, ** represents p ≤ 0.01, *** represents p ≤ 0.001 and **** represents p ≤ 0.0001.

To assess the roles of p53 and p21 in Hypo-induced cell cycle arrest, p53 or p21 genes was knocked out in RPE-1 cells via CRISPR. As expected, p53^-/-^ cells had significantly reduced expression of p21 (Figure 4B). Conversely, due to the negative feedback between p21 and p53^35^, p21^-/-^ cells had more p53 compared to wild-type (WT) cells and retained the ability to activate p53 in response to DNA damage (Figure 4B). Although p53^-/-^ and p21^-/-^ cells exhibited slower growth in Hypo compared to Medium or Iso like WT cells, p53^-/-^ and p21^-/-^ cells grew faster than WT cells in Hypo (Figure 4C and Supplementary Figure 4C). Furthermore, p53^-/-^ and p21^-/-^ cells retained significantly more S/G_2_ cells compared to WT cells after Hypo treatment and were comparable to their Iso counterparts after 72 hr (Figure 4D and Supplementary Figure 4D). The cell number and S/G_2_ population of p53^-/-^ and p21^-/-^ cells remained significantly higher than WT cells even after 7 days of continuous Hypo treatment (Supplementary Figure 4E), indicating that p53^-/-^ and p21^-/-^ cells were able to escape Hypo-induced cell cycle arrest and continue to proliferate.

As hypo-osmotic stress compromises mitotic fidelity (Figure 1), continued proliferation in the absence of p53 activation under such stress may result in aneuploidy accumulation. Chromosome spreads were performed to assess chromosome numbers of p53^-/-^ and p21^-/-^ cells after 72 hr of osmotic treatment. p53^-/-^ and p21^-/-^ cells maintained a high level of euploidy at the expected chromosome number of 46 in both Medium and Iso, but their chromosome numbers showed greater deviation from the euploid number under Hypo (Figure 4E, p = 0.0005 for p53^-/-^ cells and p = 0.002 for p21^-/-^ cells as assessed via chi-square tests). This result suggests that activation of the p53-p21 pathway to trigger G_1_ cell cycle arrest is a protective mechanism against continued error-prone mitosis that leads to genome instability (Figure 4F).

### Hypo-osmotic stress increases p53 nuclear concentration by decreasing nuclear size

Previous work showed that aneuploid RPE-1 cells can lead to p53 activation and G_1_ cell cycle arrest^36–38^. However, considering that only 30% of mitoses were erroneous compared to the near uniform cell cycle arrest under Hypo (Figure 1B and Figure 2E), aneuploidy was unlikely to be the major cause of p53 activation. Another potential inducer of p53 activation under Hypo was DNA damage^31^. However, Hypo cells did not exhibit increased accumulation of nuclear γ-H2AX foci (Supplementary Figure 5A), and only exhibited a marginal increase in total cellular p53 (Figure 4A). This minimal increase in p53 protein level is inconsistent with the robust protein stabilization that typically follows DNA damage kinase activation, which is required for strong p21 induction.

To elucidate the mechanism of p53 activation under hypo-osmotic stress, immunofluorescence was performed to assess how the endogenous levels and localizations of p53 and p21 changed after 72 hr of osmotic treatment (Figure 5A). As a positive control, 24 hr of 5 µM Nutlin (inhibiting Mdm2-mediated p53 degradation) markedly elevated both p53 and p21 levels. Although the mean p53 nuclear intensity was significantly higher in Hypo cells, the total level of nuclear p53 remained unchanged (Figure 5C). This was likely because the nuclear size of Hypo cells, as measured by the projected nuclear area, which correlated with nuclear volume (Supplementary Figure 5B), was significantly smaller than those in Medium and Iso (Figure 5B). However, p21 levels, both in terms of mean and total nuclear intensities, were significantly higher in Hypo cells (Figure 5D), consistent with increased p21 transcription and translation driven by p53 activation. These observations led us to speculate that hypo-osmotic stress does not stabilize p53 by preventing Mdm2-mediated degradation but rather concentrates p53 within smaller nuclei to drive downstream gene expression.

**Figure 5:**
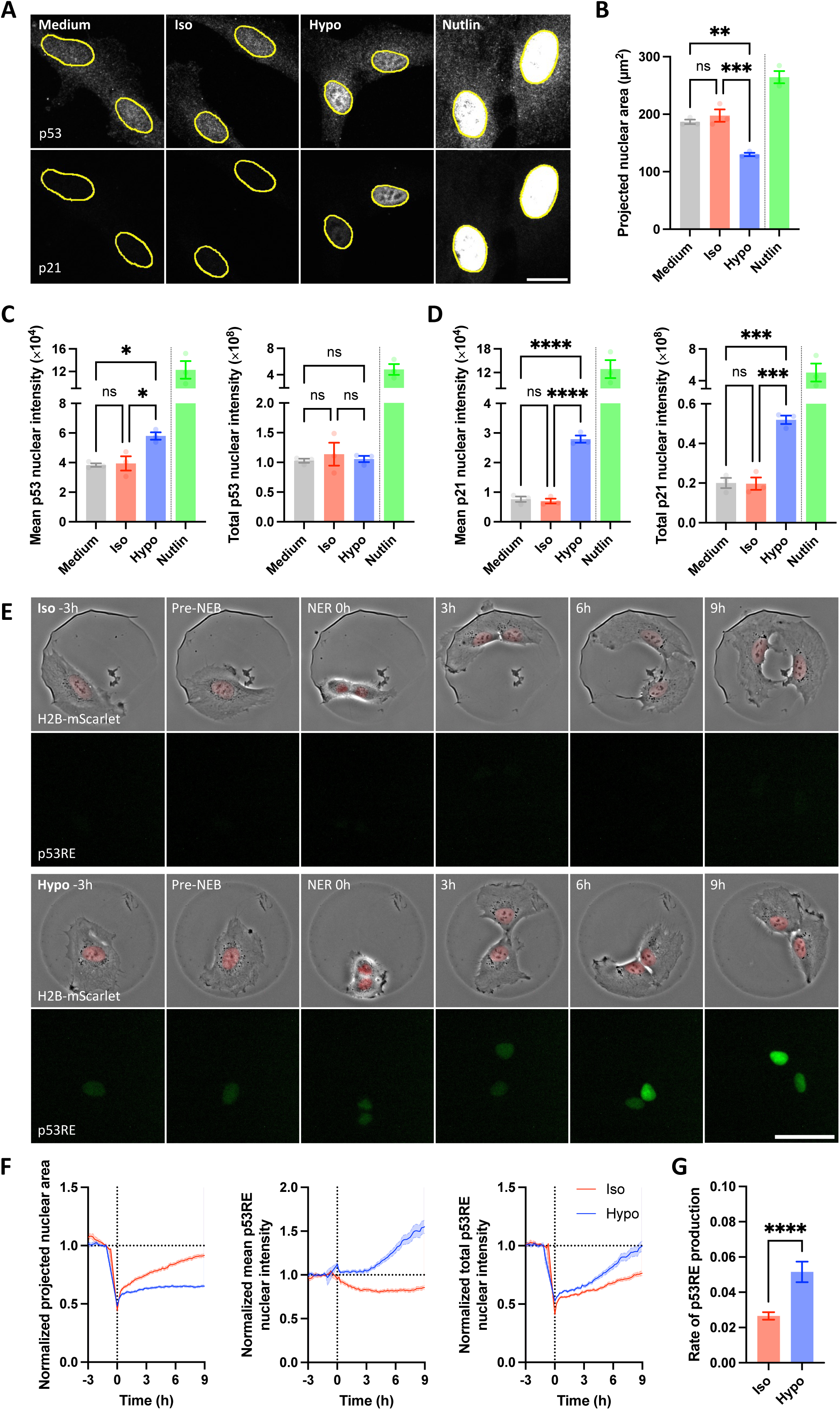
Hypo-osmotic stress activates p53 in post-mitotic daughter cells with impaired nuclear expansion. **(A)** Immunofluorescence confocal images of p53 (top) and p21 (bottom) in RPE-1 cells after 72 hr of osmotic stress. Treatment with 5 µM Nutlin for 24 hr served as a positive control to induce p53 and p21 expressions by inhibiting Mdm2-mediated p53 degradation. Nuclei were counterstained with DAPI (not shown) and outlined in yellow. Images represent z-projections from summed slices with matched intensity scaling for each channel across conditions. Nutlin-treated samples were displayed with the same scaling, resulting in image over-saturation. Scale bar = 20 µm. **(B)** Quantification of projected nuclear area. **(C)** Quantifications of mean (left) and total (right) p53 nuclear intensity. **(D)** Quantifications of mean (left) and total (right) p21 nuclear intensity. **(E)** Time-lapse montages of RPE-1 cells co-expressing H2B-mScarlet and a p53 activity reporter (p53RE) on fibronectin-coated micropatterns undergoing cell division under Iso (top) or Hypo (bottom) conditions. The montages were aligned to NER at 0 hr, and covered 3 hr before and 9 hr after NER. Scale bar = 50 µm. **(F)** Quantifications of normalized projected nuclear area (left), and normalized mean (middle) and total (right) p53RE nuclear intensity over time. The data were normalized to the pre-mitotic levels for each cell (average within 30 min prior to NEB). The graphs were aligned to NER at 0 hr, and spanned 3 hr before and 9 hr after the first mitosis. A total of 65 Gen 1 Iso cells divided to form 130 Gen 2 daughter cells, while 67 Gen 1 Hypo cells divided to form 134 Gen 2 daughter cells across four independent experiments. Line represents mean while shading represents SEM. **(G)** Quantification of the rate of p53RE production, calculated by linear fitting of normalized total p53RE nuclear intensity between 1 and 9 hr post-mitosis for each cell. All bar graphs represent mean ± SEM from n ≥ 3 independent experiments. For (B), (C) and (D), data of multiple cells from each independent experiment were averaged to generate a single biological replicate value. Statistical comparisons for (B), (C) and (D) were assessed between different osmotic conditions (excluding the positive control) with one-way ANOVA, and via t-tests for (G). ns represents not significant, * represents p ≤ 0.05, ** represents p ≤ 0.01, *** represents p ≤ 0.001 and **** represents p ≤ 0.0001.

To further examine the relationship between p53 protein accumulation and changes in nuclear size, we utilized a previously established CRISPR knock-in RPE-1 cell line where the endogenous p53 is tagged with mNeonGreen (p53-mNG)^39^. As a positive control, these cells increased their total fluorescence intensity in response to Nutlin (Supplementary Figure 5C). However, no significant difference in p53-mNG intensity was detected after 72 hr of osmotic stress treatment. Furthermore, we again observed Hypo cells to display p53-mNG enrichment, measured as mean nuclear fluorescence intensity, in smaller nuclei (Supplementary Figure 5D). Time-course analysis revealed that nuclear enrichment of p53-mNG in Hypo cells coincided with nuclear size reduction after 24 hr (Supplementary Figure 5E). The total level of nuclear p53-mNG remained comparable across all time points and treatments, further supporting that hypo-osmotic stress concentrates p53 without altering its absolute abundance.

### Hypo-osmotic stress activates p53 in post-mitotic daughter cells with impaired nuclear expansion

The 24 hr delay in nuclear size reduction and p53 nuclear enrichment (Supplementary Figure 5E) were consistent with earlier observations of time-dependent and progressive p53-mediated transcriptomic changes (Figure 3A) and cell cycle arrest (Figure 2E). To investigate the temporal pattern of Hypo-induced p53 activation, we developed a fluorescence-based p53 activity reporter, termed p53RE. It was designed based on the p53 response element of p21 promoter, placed upstream of a nuclear-targeted mNeonGreen fluorescent protein and a C-terminal EE degron (Supplementary Figure 5F). The sensitivity and specificity of p53RE were first evaluated via flow cytometry following treatment with Nutlin in WT, p53^-/-^ and p21^-/-^ cells (Supplementary Figure 5F). The mean fluorescence intensities of p53RE-expressing RPE-1 cells were assayed via flow cytometry after exposure to different osmotic stress over time. The p53 activity of Hypo-treated cells gradually increased over time from 24 to 48 hr and only became significantly higher than that of cells in Iso or Medium at 72 hr (Supplementary Figure 5G). This is in line with the previous observation that Hypo-induced cell cycle arrest only occurs after at least one round of cell division (Figure 2E), which typically happens within 24 hr, based on the average doubling time of RPE-1 cells (15-20 hr).

To correlate p53 activation and cell division, RPE-1 cells co-expressing H2B-mScarlet and p53RE were seeded on micropatterns and subjected to time-lapse imaging under Iso or Hypo conditions for 72 hr (Movie 4-5, Figure 5E). As Iso cells entered mitosis, NEB was accompanied by rapid chromosome condensation, causing a sharp decline in projected chromosomal area (Figure 5F). Two daughter cells were produced after the first mitosis, with their nuclear size and total nuclear p53RE intensity measuring at about half the pre-mitotic levels. During the subsequent interphase, nuclear size in Iso cells gradually increased back to the pre-mitotic level, along with proportional synthesis of p53RE, with the mean p53RE nuclear intensity remained mostly constant and close to baseline (Figure 5F). In contrast, Hypo cells exhibited elevated mean p53RE nuclear intensity post-mitosis, accompanied by impaired nuclear expansion (Figure 5F). Hypo daughter nuclei remained consistently smaller and never approached the pre-mitotic level. The production rate of p53RE in Hypo daughter cells surpassed that of Iso (Figure 5F-G). These data suggest that p53 activation is initiated and sustained in post-mitotic cells that fail to expand their nuclei under hypo-osmotic stress.

### p53 is activated by nuclear size reduction through a concentration effect

We postulated that limited post-mitotic nuclear expansion under Hypo condition increases the nuclear concentration of p53, and that this enrichment, rather than a major change in total protein stability and abundance of p53, is sufficient to trigger downstream transcriptional activation and cell cycle arrest. Consistent with this hypothesis, only post-mitotic Hypo cells, which experienced pronounced nuclear size reduction, exhibited robust p53 activation, whereas pre-mitotic and non-mitotic Hypo cells displayed minimal change in nuclear size and much milder p53RE expression (Supplementary Movie 1, Supplementary Figure 6A-B). Importantly, completion of mitotic cell division achieves a sufficiently small nuclear volume (i.e. approximately half of the pre-mitotic size) in the daughter cells to enable nuclear p53 concentration to surpass its activation threshold.

To theoretically evaluate whether limited post-mitotic nuclear expansion under hypo-osmotic stress was sufficient to drive the observed p53 activation, we developed a simple ordinary differential equation (ODE)-based model (Figure 6A) to describe the dynamics of the normalized total p53RE reporter, where the synthesis is driven by the nuclear concentration of p53 (total nuclear p53/nuclear volume). This model incorporated the key assumption of comparable total nuclear p53 levels that does not vary with stress, as indicated by numerous lines of evidence described earlier (Figure 5C, Supplementary Figure 5C-E). Indeed, we observed similar p53 protein dynamics across osmotic conditions, as demonstrated by negligible differences in the rates of p53 synthesis, degradation and nuclear transport (Supplementary Figure 6C-E). Applying this key assumption reduces the system to a single equation where the only differential factor between Iso and Hypo conditions is the time-dependent post-mitotic nuclear size changes (see Methods and Supplementary Information for more details regarding model development).

**Figure 6:**
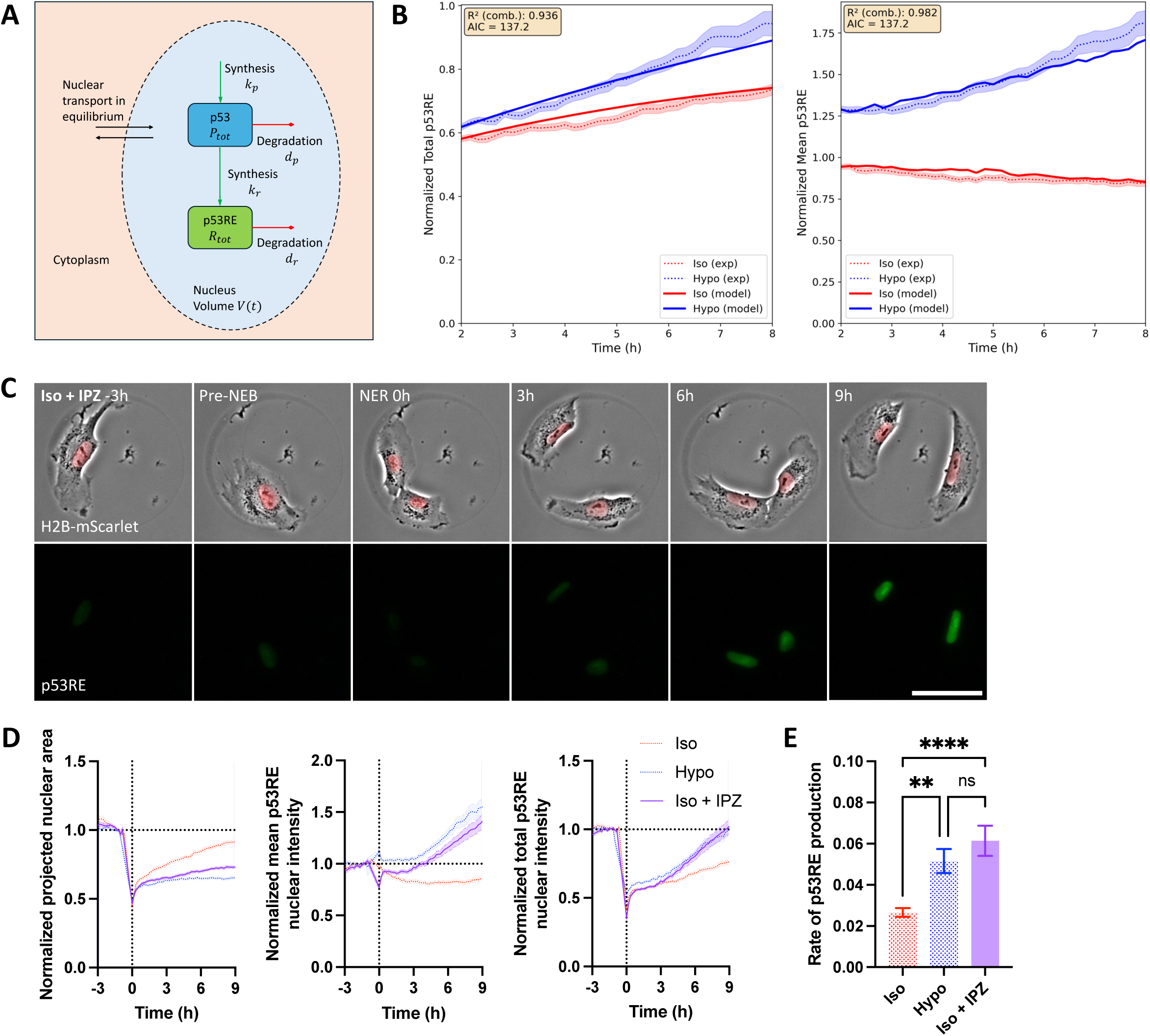
p53 is activated by nuclear size reduction through a concentration effect. **(A)** Schematic of the proposed mathematical model of p53RE dynamics. 𝑃_𝑡𝑜𝑡_ represents the total amount of nuclear p53, while 𝑅_𝑡𝑜𝑡_ represents the total amount of nuclear p53RE. 𝑉(𝑡) represents nuclear volume. 𝑘_𝑝_ and 𝑘_𝑟_ are the synthesis rate constants, and 𝑑_𝑝_and 𝑑_𝑟_are the degradation rate constant for p53 and p53RE respectively. **(B)** Model-predicted normalized total p53RE (left) and p53RE concentration (right) over time, compared with the experimental data from Figure 5F. Best-fit parameters: p53RE effective synthesis rate 𝑘_𝑠𝑦𝑛,𝑒𝑓𝑓_ = 0.0388, p53RE degradation rate 𝑑_𝑟_= 0.0385. Model-predicted and experimental p53RE concentrations were calculated by simply dividing the respective total p53RE values by the mean experimental nuclear volume data (all normalized). Dotted lines represent experimental data with SEM shown as shaded areas, while solid lines represent the model fit. Akaike Information Criterion (AIC) and R^2^ values are indicated. **(C)** Time-lapse montages of RPE-1 cells co-expressing H2B-mScarlet and p53RE on fibronectin-coated micropatterns undergoing cell division under Iso condition with 10 µM Importazole (IPZ). The montages were aligned to NER at 0 hr, and covered 3 hr before and 9 hr after NER. Scale bar = 50 µm. **(D)** Quantifications of normalized projected nuclear area (left), and normalized mean (middle) and total (right) p53RE nuclear intensity over time. The data were normalized to the pre-mitotic levels for each cell (average within 30 min prior to NEB). The graphs were aligned to NER at 0 hr and spanned 3 hr before and 9 hr after the first mitosis. A total of 56 Gen 1 Iso + IPZ cells divided to form 112 Gen 2 daughter cells across four independent experiments. The data for Iso- and Hypo-treated cells from Figure 5F were superimposed for ease of comparison. Line represents mean while shading represents SEM. **(E)** Quantification of the rate of p53RE production, calculated by linear fitting of normalized total p53RE nuclear intensity between 1 and 9 hr post-mitosis for each cell. The data for Iso- and Hypo-treated cells from Figure 5G were also included for comparison with Iso + IPZ with one-way ANOVA. Bar graph in (E) represents mean ± SEM from n ≥ 3 independent experiments. Statistical comparisons for (E) were with one-way ANOVA. ns represents not significant, * represents p ≤ 0.05, ** represents p ≤ 0.01, *** represents p ≤ 0.001 and **** represents p ≤ 0.0001.

When compared to the experimental data (Figure 5F), the model accurately recapitulated the experimental p53RE dynamics observed under both Iso and Hypo conditions (Figure 6B). It successfully captured the elevated rate of total p53RE accumulation under Hypo relative to Iso conditions, achieving a high combined coefficient of determination (R² = 0.936). Furthermore, the model’s predictions for mean p53RE, obtained by dividing the predicted total p53RE by the experimental volume (estimated from the projected nuclear surface area, see Methods for details), also showed excellent agreement (R² = 0.982). These results demonstrate that a simple concentration-driven mechanism, in which p53RE synthesis scales with nuclear p53 concentration and nuclear volume dynamics, is sufficient to explain the differential p53 activation within the 2-8 hr post-mitotic window. The choice of this simple model was further corroborated in Supplementary Information, where a formal statistical comparison against more complex model variants that incorporated non-linear volume scaling and/or condition-specific synthesis rates showed only marginal improvements in fit.

To experimentally test the effect of failed post-mitotic nuclear expansion on p53 activation, cells were treated with 10 µM Importazole (IPZ) under Iso condition and subjected to the same p53 activity measurement (Movie 6, Figure 6C). IPZ inhibits importin-β, resulting in decreased nuclear import and nuclear size (Supplementary Figure 6F). At the moderate concentration of 10 µM, IPZ did not interfere with mitotic entry or completion, but nearly all IPZ-treated cells ceased division following the first mitosis, phenocopying the Hypo-induced post-mitotic cell cycle arrest (Figure 6C). Notably, nuclei of IPZ-treated cells under Iso stopped expanding and remained consistently small post-mitosis, leading to similar increase in mean p53RE nuclear intensity comparable to that of Hypo-treated cells (Figure 6D). The production rate of p53RE in IPZ-treated daughter cells also mirrored that of Hypo and was significantly higher than that of untreated Iso control (Figure 6D-E). Together, these findings demonstrate that nuclear size reduction alone was sufficient to activate p53 through a concentration-driven mechanism.

### Altered chromatin compaction contributes to impaired nuclear expansion and p53 activation

Nuclear growth at mitotic exit is a multi-step process that requires coordinated chromatin decondensation, nuclear envelope reassembly and active nucleocytoplasmic transport^40,41^. Previous study^42^ and our data suggest that nucleocytoplasmic transport was largely unaffected by hypo-osmotic stress (Supplementary Figure 6D-F). Instead, nuclei in cells under Hypo exhibited a more heterogenous Hoechst staining pattern with bright condensed chromatin foci than those under Iso (Figure 7A). This heterogeneity, quantified as the coefficient of variation (COV) of the Hoechst signal, was significantly higher in Hypo cells (Figure 7B). Moreover, cells under Hypo possessed lower level of H4K16ac, an epigenetic histone modification that marks decondensed chromatin, compared to Iso, as well as

**Figure 7:**
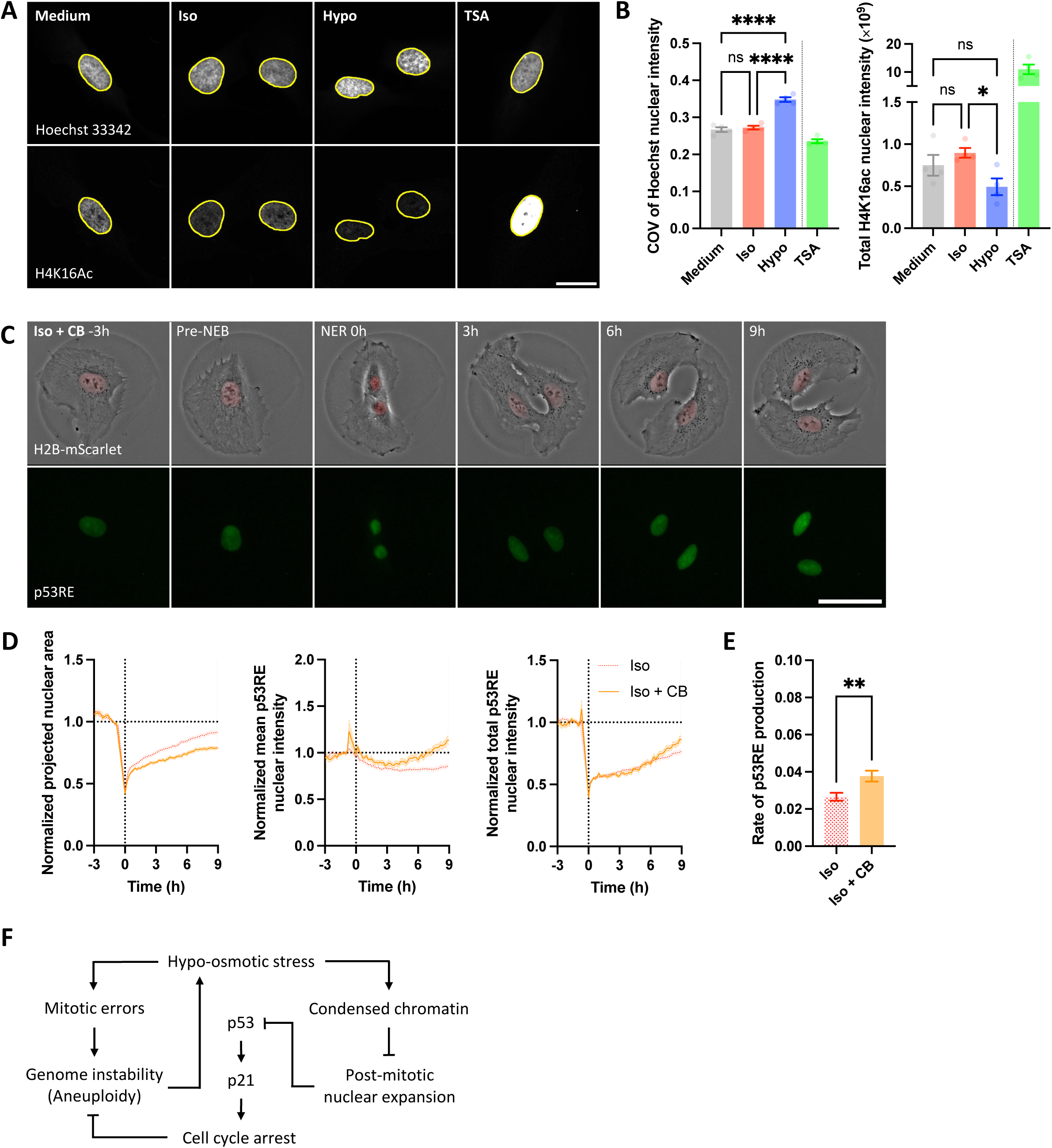
Altered chromatin compaction contributes to impaired nuclear expansion and p53 activation. **(A)** Immunofluorescence confocal images of nuclei/chromatin (Hoechst 33342, outlined in yellow, top) and H4K14Ac (bottom) in RPE-1 cells after 72 hr of osmotic stress. Treatment with 100ng/mL Trichostatin A (TSA) for 24 hr served as a positive control to decondense chromatin. Images represent z-projections from summed slices with matched intensity scaling for each channel across conditions. TSA-treated samples were displayed with the same scaling, resulting in image over-saturation for H4K14Ac. Scale bar = 20 µm. **(B)** Quantifications of chromatin heterogeneity by coefficient of variation (COV) of Hoechst nuclear intensity (left), and total H4K16ac nuclear intensity (right). **(C)** Time-lapse montages of RPE-1 cells co-expressing H2B-mScarlet and p53RE on fibronectin-coated micropatterns undergoing cell division under Iso condition with 0.5µM CB-6644 (CB). The montages were aligned to NER at 0 hr, and covered 3 hr before and 9 hr after NER. Scale bar = 50 µm. **(D)** Quantifications of normalized projected nuclear area (left), and normalized mean (middle) and total (right) p53RE nuclear intensity over time. The data were normalized to the pre-mitotic levels for each cell (average within 30 min prior to NEB). The graphs were aligned to NER at 0 hr and spanned 3 hr before and 9 hr after the first mitosis. A total of 53 Gen 1 Iso + CB cells divided to form 106 Gen 2 daughter cells across three independent experiments. The data for Iso-treated cells from Figure 5F were superimposed for ease of comparison. Line represents mean while shading represents SEM. **(E)** Quantification of the rate of p53RE production, calculated by linear fitting of normalized total p53RE nuclear intensity between 1 and 9 hr post-mitosis for each cell. The data for Iso-treated cells from Figure 5G were also included for comparison with Iso + CB with t-tests. **(F)** Schematic summary of the proposed mechanism of p53-activation under hypo-osmotic stress. Post-mitotic cells under hypo-osmotic stress exhibit hyper-condensed chromatin and impaired expansion of daughter nuclei, resulting in smaller nuclear volume. The spatial constraint increases the nuclear concentration of p53 without a substantial change in the total protein level of p53, leading to p53 activation and a p53/p21-dependent G_1_ cell cycle arrest that prevents further error-prone divisions. All bar graphs represent mean ± SEM from n ≥ 3 independent experiments. For (B), data of multiple cells from each independent experiment were averaged to generate a single biological replicate value. Statistical comparisons for (B) were assessed between different osmotic conditions (excluding the positive control) with one-way ANOVA. ns represents not significant, * represents p ≤ 0.05, ** represents p ≤ 0.01, *** represents p ≤ 0.001 and **** represents p ≤ 0.0001.

Trichostatin A (TSA)-treated positive control cells that accumulated H4K16ac due to histone deacetylase inhibition (Figure 7A-B). To further examine chromatin compaction, Fluorescence Lifetime Imaging - Förster Resonance Energy Transfer (FLIM-FRET) measurements were made in RPE-1 cells co-expressing H2B-mNeonGreen and H2B-mCherry^43^. Cells under Hypo displayed a significantly higher FRET (lower fluorescence lifetime) than those under Iso. The higher FRET indicated higher chromatin compaction, in contrast to lower FRET (higher fluorescence lifetime) and less condensed chromatin in TSA-treated cells (Supplementary Figure 7A).

To test whether impaired nuclear expansion due to lack of chromosome decondensation can cause p53 activation, CB-6644 (CB) was used to inhibit the RUVBL1/2 complex known to be critical for chromatin decondensation at mitotic exit^44,45^. CB treatment in Iso cells led to a significant reduction in nuclear size with more heterogenous Hoechst staining (Supplementary Figure 7B). This was accompanied by increased mean nuclear p53-mNG intensity without significantly altered total intensity (Supplementary Figure 7C). Live imaging was performed to monitor nuclear dynamics and p53 activation with or without CB treatment under Iso condition (Movie 7, Figure 7C). CB-treated cells exhibited similar compromised post-mitotic nuclear growth as Hypo or IPZ treatment, though to a milder degree, resulting in a modest increase in mean p53 nuclear intensity and p53RE production rate that were higher than untreated daughter cells (Figure 7D-E). These data suggest that a defect in chromatin decondensation of post-mitotic nuclei under Hypo could be sufficient to impede nuclear expansion and activate p53.

## Discussion

The results presented above demonstrate that despite swift initial volume recovery, sustained hypo-osmotic stress, likely through profound alteration in intracellular milieu, elicits a series of effects on the cell that delay mitosis and compromise chromosome segregation fidelity. Importantly, hypo-osmotic stress also triggers a p53/p21-dependent G_1_ cell cycle arrest that suppresses further error-prone divisions and chromosome instability (Figure 7F). This p53 response is distinct from the canonical mechanism of p53 activation that depends on post-translational modifications of p53 by DNA damage sensor kinases (i.e. ATM and ATR), which decrease the interaction of p53 with Mdm2 and subsequent degradation^32^. Under hypo-osmotic stress, however, we detected neither increased DNA damage nor robust p53 stabilization and protein level increase. Instead, p53 is primarily induced in post-mitotic daughter cells with impaired nuclear expansion, where a smaller nuclear compartment is sufficient to passively elevate nuclear p53 concentration beyond its transactivation threshold to drive downstream gene expression without requiring substantial changes in total p53 abundance (Figure 7F).

This concentration-dependent mode of transcriptional activation is reminiscent of the developmental timer that regulates the onset of zygotic genome activation^46^. During early embryonic development, repeated mitotic cleavage divisions rapidly increase the number of nuclei without growth and proportional increase in cytoplasmic volume. The resultant rise in the ratios of genetic material or nuclear volume to cytoplasm effectively dilutes maternally inherited transcriptional repressors and titrates away their inhibitory effects, in turn permitting the engagement of pioneer transcription factors such as DUX4 to open compacted zygotic chromatin and increase their accessibility for transcription. Progressive nuclear size reduction from repeated mitoses also serves to concentrate transcription factors and chromatin modifiers above their critical activation concentrations^47^. We propose that similar concentration-driven activation model may represent a broader and evolutionarily conserved principle by which cells modulate transcriptional outputs through nuclear size-sensing.

Although our results suggest that nuclear size reduction is sufficient and likely a major contributor towards p53 activation, we cannot completely exclude auxiliary biochemical contributions. Besides the canonical stress-activated kinases (i.e. ATR, ATM and CHK1/2) that phosphorylate either p53 or Mdm2 to alter the stability and degradation kinetics of p53, p53 can also be post-translationally modified, most notably with acetylation and methylation, to enhance its nuclear retention, DNA binding and transactivation ability^48,49^. Additionally, p53 can also be activated transcriptionally^50^. Our RNA-seq data reveals a modest time-dependent but statistically insignificant trend in *TP53* mRNA upregulation under hypo-osmotic stress (Figure 3B Cluster 4, Supplementary Figure 4A). These parallel biochemical mechanisms may still cooperatively augment p53 activation when nuclear expansion is restricted.

Recent studies defined a mechanism of p53 activation, termed mitotic stopwatch, which proposes that prolonged mitotic arrest exceeding 90 min stabilizes and activates p53 through accumulation of mitotic stopwatch complex components consisting of 53BP1, USP28, Plk1 and p53^51–53^. In cases of moderate mitotic delay between 30 to 90 min, the complex, at sub-threshold level, can be retained and transmitted to following generations, and arrest cells if mitosis is moderately delayed again^54^. Several features distinguish the mechanism of p53 activation uncovered in this study from the mitotic stopwatch. First, the mitotic delay induced by hypo-osmotic stress (10 to 20 min) is considerably shorter than the threshold of 90 min required for mitotic stopwatch activation. Second, despite only experiencing moderate delay, most hypo-osmotic cells activate p53 and trigger arrest without requiring a second round of delayed mitosis. Third, hypo-osmotic stress activates p53 through a nuclear concentration-based mechanism without increasing the total protein level of p53. Lastly, hypo-osmotic stress-induced cell cycle arrest is reversible upon osmotic normalization, indicating a cytoprotective adaptation rather than an irreversible senescent outcome characteristic of mitotic stopwatch-mediated arrest.

The exact mechanism by which hypo-osmotic stress inhibits post-mitotic nuclear expansion is still an open question. We propose that a reduction in intracellular ionic strength following equilibration to external hypo-osmotic environment may disfavor chromatin decondensation. It has been proposed that chromatin condensation releases while decondensation absorbs counterions^55^. It is possible that hypo-osmotic conditions with a decreased intracellular ion reserve favor a more condensed chromatin state. As the nuclear envelope is physically tethered to the chromatin surface via inner nuclear membrane proteins^56^, this condensed state would inherently resist expansion, providing a plausible biophysical basis for the observed reduction in nuclear size. Notably, this chronic stress-induced hyper-condensation is distinct from the chromatin decondensation observed during acute hypo-osmotic stress^57–59^. In these studies, cells were typically treated transiently and observed for short periods of time, ranging from 15 min to 3 hr. Importantly, although chromatin remained decondensed for several hours, they subsequently recovered and became hyper-condensed after 24 hr of sustained hypo-osmotic stress^60^, consistent with our observations. These stress duration-dependent changes in chromatin condensation state highlight the complexity of chromatin organization *in vivo*, which is influenced by dynamic processes such as macromolecular crowding and chromatin remodeling as cells acclimatize to osmotic imbalance and actively adapt to changes in intracellular ionic composition.

Another observation worthy of further investigation is the increased mitotic error rate under hypo-osmotic stress. Changes in cytoplasmic viscosity, ionic composition and macromolecular crowding following regulatory volume decrease could potentially affect the phase separation behaviors of mitotic structures. For example, pericentriolar materials are known to phase-separate and self-assemble into distinct compartment to concentrate, organize and nucleate tubulin to form mitotic spindles^61^. Phase separation of DNA binding protein Ki-67 had been shown to modulate the steric hindrance and electrostatic interactions between chromosomes and aid in their proper dispersal during anaphase^62^. Chromosome passenger complexes are also known to phase-separate around inner centromeres and influence the dynamics of chromosome attachment and segregation^63^. Perturbations of these condensates may underlie defects in spindle assembly, chromosome organization and microtubule attachment, as suggested by signs of activation of the spindle assembly checkpoint.

Finally, p53^-/-^ and p21^-/-^ cells do not arrest but still proliferate slower under hypo-osmotic stress, suggesting that their cell cycles are generally extended. Extended interphase is reflective of slower protein synthesis and biomass accumulation. Our RNA-seq results also identify consistent and progressive changes in sugar metabolism and glycosylation (Figure 3C) which may lead to altered nutrient utilization and cellular energetics in cells growing in hypo-osmotic environment. Recent studies show that external mechanical stress, including hypo-osmotic stress, induces disassembly of membrane caveolae, releasing caveolar components that can participate in downstream signaling^64–66^. Additionally, caveolae disassembly can also potentially hamper endocytosis^67^. This is in line with our previous observation that aneuploid cells, which experience hypo-osmotic like stress, likewise exhibit endocytic defects ^25^.

In conclusion, we demonstrate that hypo-osmotic stress impairs mitotic fidelity and triggers a mechanosensitive p53-dependent cell cycle arrest to avert error-prone mitosis and preserve genome stability. This concentration-driven p53 activation establishes nuclear size as a key and previously underappreciated determinant of stress sensing and proliferation control. We propose that this mechanism of p53 regulation is critical for maintaining tissue homeostasis and preventing osmotic stress-induce chromosome instability that can lead to expanded genetic and phenotypic diversity that predispose diseases such as inflammation-associated cancer.

## Materials and Methods

### Mammalian cell culture

Cell culture was performed in accordance with American Type Culture Collection guidelines, with minor modifications. hTERT RPE-1 cells were maintained in Dulbecco’s Modified Eagle Medium/Nutrient Mixture F-12 (DMEM/F12, Gibco, 11320033) supplemented with 10% (v/v) heat-inactivated fetal bovine serum (FBS, Gibco, 16140071) and 100 units/mL penicillin-streptomycin (P/S, Gibco, 15140122). BJ-5ta and MDCK cells were cultured in DMEM (Gibco, 11965092) supplemented with 10% (v/v) FBS and 100 units/mL P/S. Nalm6 cells were maintained in RPMI 1640 Medium (Gibco, 11875093) with the same supplements. Mouse colonoids were isolated and maintained as previously described^36^, using conditioned medium collected from the L-WRN cell line following an establish protocol^68^. All cells were cultured at 37°C in a humidified incubator with 5% CO_2_.

### Lentivirus production

HEK-293T cells were utilized to produce lentiviral particles. The cells were maintained in DMEM supplemented with 10% (v/v) FBS and 100 units/mL P/S. To produce lentivirus, HEK-293T cells were co-transfected in antibiotic-free medium using polyethylenimine (Polysciences, 24765) at a polymer-to-DNA ratio of 3:1 with the plasmid of interest and third-generation lentiviral packaging plasmids, comprising pMDLg/pRRE (Addgene, 12251), pRSV-Rev (Addgene, 12253) and pMD2.G (Addgene, 12259). Cell culture medium was replaced after 24 hr, and conditioned medium containing lentivirus was collected at 48 and 72 hr post-transfection. The virus-containing conditioned medium was filtered through a 0.45 µm syringe filter (Sartorius, S6555) and stored at −80°C. Lentiviruses encoding FUCCI cell cycle marker (mAG-hGeminin and mKo2-SLBP), H2B-mNeonGreen, H2B-mCherry, H2B-mScarlet, p53RE and NLS-GFP were produced and used in this study.

### Cell line generation

Fluorescently labeled RPE-1 cells expressing FUCCI, H2B-mNeonGreen, H2B-mCherry, H2B-mScarlet, p53RE and NLS-GFP were generated by lentiviral transduction. Wild-type (WT) RPE-1 cells were incubated with conditioned medium containing the respective lentivirus for 24 to 72 hr, followed by fluorescence-activated single-cell sorting into 96-well plates with a Sony SH800 cell sorter. Clonal populations exhibiting optimal expression patterns and intensities were expanded. p53^-/-^ and p21^-/-^ RPE-1 cells were generated by CRISPR-Cas9-mediated genome editing. WT cells were co-electroporated with Cas9 nuclease (IDT, 1081059) and a guide RNA duplex comprising ATTO 550 tracrRNA (IDT, 1075928) and target-specific crRNA using the Neon Transfection System (Invitrogen). Two crRNAs were used for each gene target: Hs.Cas9.TP53.1.AA (IDT, 110796316) and Hs.Cas9.TP53.1.AC (IDT, 110796317) for p53, and Hs.Cas9.CDKN1A.1.AA (IDT, 110796318) and Hs.Cas9.CDKN1A.1.AB (IDT, 110796319) for p21. Transfected (FL2-positive) cells were single-cell sorted into 96-well plates 24 hr post-electroporation and expanded for two weeks. Putative knockout clones were initially screened for Nutlin resistance, and successful knockout was confirmed by Western blot analysis of p53 and p21 protein expression.

### Osmotic stress and drug treatment

Hypo-osmotic cell culture media of decreasing osmolality were formulated by diluting standard culture medium with progressively higher proportions of deionized water. Corresponding isotonic controls were also prepared to account for nutrient dilution by diluting the same culture medium with Dulbecco’s PBS without calcium and magnesium (Gibco, 14190144) at the same proportions. All media were sterile-filtered through 0.2 µm syringe filters (Sartorius, S6534), and their osmolarities were measured by freezing point depression using an Advanced Instruments Micro-Osmometer (Precision System, OSMETTE, Model 5004). In selected experiments, cells were incubated with drugs diluted in the respective osmotic media for 24 to 72 hr. The following pharmacological inhibitors were used: Etoposide (Sigma-Aldrich, E1383, 10 µM), Nutlin (Sigma-Aldrich, N6287, 5 µM), Doxorubicin hydrochloride (Sigma-Aldrich, D1515, 1 µM), Importazole (Selleck Chemicals, S8446, 10 µM), Trichostatin A (MedChem Express, HY-15144, 100ng/mL) and CB-6644 (MedChem Express, HY-114429, 0.5µM).

### Cell proliferation assay

Cells were seeded at 5×10^4^ per well in 6-well plates with standard cell culture medium and allowed to adhere overnight (−24h). Medium was replaced with osmotic stress media the next day (0h), and cells were maintained in these conditions for 72 hr without media replacement. At 24 hr intervals (24, 48 and 72h), cells were harvested with 0.25% trypsin-EDTA (Gibco, 25200056) and resuspended in 100µL of medium. Prior to counting, cell morphology was documented by brightfield imaging at 4× magnification using an EVOS M5000 Imaging System (Invitrogen). Viable cell numbers were quantified with a LUNA-II Automated Cell Counter (Logos Biosystems) following exclusion of dead cells by 0.4% trypan blue stain (Logos Biosystems, T13001). Growth rates were calculated by fitting data from at least three biological replicates to a standard exponential growth model using GraphPad Prism.

### Flow cytometry

Cell cycle distribution was assessed by DNA content analysis following osmotic stress. Cells were harvested by trypsinization, collected in 1.5mL microcentrifuge tubes, and stained with Hoechst 33342 (Sigma-Aldrich, SC200908, 1µg/mL) diluted in medium for 45 min at 37°C with periodic gentle mixing. Stained cells were analyzed with a Sony SH800 cell sorter to determine the proportion of cells in different cell cycle phases based on DNA content. For fluorescence quantification of p53RE or p53-mNG expression, osmotically treated reporter cells were collected and analyzed directly with a Sony SH800 cell sorter without fixation or additional staining. Fluorescence intensity distributions were plotted as histograms using FlowJo, and mean fluorescence intensities were quantified and compared. All flow cytometry experiments included a minimum of 10000 events per sample and were performed in at least three independent biological replicates.

### Micropattern fabrication

Fibronectin-coated circular micropatterns were fabricated using a modified soft lithography technique based on a published protocol^69^. Polydimethylsiloxane (PDMS) molds were first made by pouring a 10:1 mixture of PDMS base and crosslinker onto a silicon wafer with an array of cylindrical features of 8000 µm^2^ in cross-sectional area. This size was selected based on the measured projected area of RPE-1 cells (1500-2000 µm^2^), which allows one cell to undergo two complete rounds of cell division without experiencing crowding and contact inhibition. After curing at 80°C for 2 hr, PDMS stamps were cut into appropriate size and treated with oxygen plasma for 90 sec (Harrick Plasma Cleaner), before being placed pattern-side down on a 35 mm glass-bottom uncoated dish (Ibidi, 81151). 5µL of NOA 63 (Edmund Optics, #16-778) was introduced into the spacing between the PDMS mold and glass substrate, followed by UV polymerization (UV-KUB 2, KLOE) at 20% power for 20 sec to create a transparent protective mask. After PDMS mold removal, the dish was coated with 50 µg/mL fibronectin (Roche, 10838039001) overnight at 4°C to functionalize the exposed glass areas with extracellular matrix for cell adhesion. Following PBS washing, the NOA 63 mask was removed, and the dish was passivated with 0.2% (w/w) Pluronic F-127 (Sigma-Aldrich, P2443) for 30 min to block non-patterned regions. Completed micropatterned dishes were stored in PBS at 4°C for up to one month.

### Cell seeding on micropatterns

RPE-1 cells were suspended in standard cell culture medium at a density of 1×10⁶ cells/mL. 100µL of cell suspension was dispensed onto the micropatterned region and incubated at 37°C for 2 min to facilitate cell attachment. After 2 min, unbound cells were removed by gentle aspiration, and the micropatterned surface was rinsed twice with fresh medium. Seeding efficiency was verified under a bright-field microscope, with the goal of achieving single-cell occupancy for majority of the array. Successfully seeded dishes were supplemented with 2mL of medium and incubated at 37°C for 2 hr to allow for cell spreading before live-cell imaging.

### Time-lapse live-cell imaging

RPE-1 cells expressing FUCCI or H2B-mScarlet/p53RE reporters were seeded on micropatterned dishes as described above. Medium was replaced with experimental osmotic media prior to imaging. Imaging was performed on a Nikon Biostation IM-Q system equipped with phase-contrast and fluorescence capabilities and a humidified environmental chamber at 37°C with 5% CO₂. For FUCCI experiments, images were acquired every 10 min for 72 hr using a 20× objective lens. For p53RE experiments, images were captured every 10 min for 72 hr using a 40× objective lens. Stage positions were recorded to enable continuous tracking of individual cells across time points. Image analyses were performed in ImageJ. H2B-mScarlet signal was used for automated nuclear segmentation and tracking with the TrackMate plugin to correlate nuclear dynamics and p53 activation relative to cell division under osmotic stress. To monitor mitosis under osmotic stress, RPE-1 cells expressing H2B-mNeonGreen were seeded on 35 mm glass-bottom dishes (Iwaki, 3970-035) and subjected to high-resolution confocal time-lapse live-imaging. Imaging was performed on a W1-Dual Cam Spinning Disk microscope with a 40× 1.3 N.A. oil immersion objective lens and a humidified chamber maintained at 37°C with 5% CO₂. Z-stacks (30-40 µm range, 1 µm steps) were acquired every 4 min for 24 hr using the Metamorph software, with stage positions recorded for longitudinal tracking of individual cells. Mitotic events were analyzed from maximum intensity projections, and errors were manually scored. AB were counted when strands of chromatin material were observed between segregating chromatin masses during anaphase. LC were counted when one or more chromosomes fail to keep pace with segregating chromatin masses during anaphase.

### Immunofluorescence

Cells were seeded on glass-bottom dishes (Iwaki, 3970-035 or 3911-035) and treated with osmotic stress with or without drugs for defined durations, before fixing with 4% paraformaldehyde in PBS (Santa Cruz Biotechnology, sc-281692) for 15 min at room temperature (RT). After PBS rinsing, samples were permeabilized with 0.3% (v/v) Triton X-100 (Sigma-Aldrich, X100) in PBS for 15 min, and blocked with 5% (w/v) bovine serum albumin (BSA, Sigma-Aldrich, A2153) in PBS containing 0.3% (v/v) Triton X-100 for 2 hr at RT. Samples were incubated with primary antibodies diluted in antibody dilution buffer, comprising PBS with 2% (w/v) BSA and 0.2% (v/v) Triton X-100, at 4°C overnight. The following primary antibodies were used in this study: anti Ki-67 (Cell Signaling Technology, 9129S, rabbit monoclonal, 1:400), anti γ-H2AX (Cell Signaling Technology, 80312S, mouse monoclonal, 1:400), anti p53 (Abcam, ab1101, mouse monoclonal, 1:200), anti p21 (Cell Signaling Technology, rabbit monoclonal, 2947S, 1:500) and anti H4K16Ac (Cell Signaling Technology, 13534S, rabbit monoclonal, 1:500). Primary antibodies were removed on the following day, and samples underwent three sequential 15 min washes with PBS containing 0.2% (v/v) Triton X-100, PBS containing 0.2% (v/v) Tween 20 (Sigma-Aldrich, P1379), and PBS alone. After the wash cycle, samples were incubated with fluorophore-conjugated secondary antibodies diluted in antibody dilution buffer for 2 hr at RT protected from light. The following secondary antibodies were used in this study: Alexa Fluor 546 goat anti-mouse IgG (Invitrogen, A11003, 1:250), Alexa Fluor 647 goat anti-rabbit IgG (Invitrogen, A27040, 1:250), and Alexa Fluor 488 goat anti-rabbit IgG (Invitrogen, A27034, 1:500). Selected samples were counterstained with Alexa Fluor 488 Phalloidin (Cell Signaling Technology, 8878S, 1:800), DAPI (Sigma-Aldrich, MBD0015, 1µg/mL) or Hoechst 33342 (Sigma-Aldrich, SC200908, 1µg/mL) diluted in PBS for 1 hr at RT protected from light. Finally, samples were washed with the same three-solution series and stored in PBS at 4°C until imaging. Fluorescence images were acquired with a Zeiss LSM980 AiryScan confocal microscope with 20× 0.8 N.A. air or 63× 1.4 N.A. oil immersion objectives. At least 10 images, each containing 1 or more cells, were taken per sample. Data from each staining experiment were averaged to generate a single biological replicate value. Each staining experiment were repeated at least three independent times. Data were visualized, processed and analyzed using ImageJ.

### EdU cell proliferation assay

Edu staining was performed using the EdU Assay/EdU Staining Proliferation iFluor 488 Kit (Abcam, ab219801) following the manufacturer’s protocol. After osmotic treatments, cells were pulse-labeled with EdU for the specified duration before fixation and counterstaining with Hoechst 33342 at 1µg/mL. Fluorescence imaging was acquired with the same imaging protocol as described earlier. At least 10 representative fields containing multiple cells were taken per sample using a 20× objective lens. The percentage of EdU-positive cells was calculated from the ratio of EdU-stained nuclei over the total number of Hoechst-stained nuclei per field of view. Data from the 10 fields were averaged to generate a single biological replicate value, and the entire assay was repeated three independent times.

### Western blot

Cells were lysed directly in culture dishes on ice using a cell scraper and an appropriate volume of RIPA lysis buffer (Thermo Scientific, 89900) supplemented with 1× Protease/Phosphatase Inhibitor Cocktail (Cell Signaling Technology, 5872S) after medium removal and PBS washing. Lysates were further homogenized by vortexing and centrifuged at 13000 rpm at 4°C for 15 min. Supernatant were collected into 1.5mL microcentrifuge tubes and stored at −80°C until further processing. Protein concentrations of the cell lysates were measured using a BCA Protein Assay Kit (Thermo Scientific, 23225) and a CLARIOstar Plus Plate Reader (BMG Labtech). Equal protein amounts (10-20µg) were denatured in 1× Reducing Laemmli SDS Sample Buffer (Thermo Scientific, AAJ61337AD) at 95°C for 10 min. Protein samples were loaded into 4-20% gradient polyacrylamide Mini-PROTEAN TGX gels (Bio-Rad, 4561094) and resolved by electrophoresis at constant 200V for 30 min. Resolved proteins were then electroblotted onto 0.2 µm nitrocellulose membranes using the Trans-Blot Turbo Transfer System (Bio-Rad, 1704158). Transfer efficiencies were verified by Ponceau S staining (Cell Signaling Technology, 59803S), before blocking with PBS-based Intercept Blocking Buffer (LI-COR 927-70001) for 2 hr at RT with gentle agitation. Membranes were probed overnight at 4°C overnight on an orbital shaker with primary antibodies diluted in antibody dilution buffer, comprising 1:1 Intercept Blocking Buffer:PBS containing 0.2% (v/v) Tween 20. The following primary antibodies were used in this study: anti p53 (Abcam, ab1101, mouse monoclonal, 1:500), anti p21 (Cell Signaling Technology, 2947S, rabbit monoclonal, 1:1000) and anti β-actin (Invitrogen, MA1-140, mouse monoclonal, 1:10000). On the following day, membranes were washed 3× for 5 min with PBS containing 0.1% (v/v) Tween 20, followed by 2× for 5 min with PBS, before incubation with secondary antibodies diluted in antibody dilution buffer for 2 hr at RT in the dark with gentle agitation. Infra-red dye-conjugated secondary antibodies were used in this study: goat anti-mouse 800CW (LI-COR, 926-32210, 1:10000), goat anti-rabbit 800CW (LI-COR, 926-32211, 1:10000), goat anti-mouse 680LT (LI-COR, 926-68020, 1:10000) and goat anti-rabbit 680LT (LI-COR, 926-68021, 1:10000). The washing steps as described earlier were repeated. Infra-red signals of the protein bands were detected using an Odyssey CLx Scanner (LI-COR) and quantified in Image Studio Lite (LI-COR). Band intensities were normalized to β-actin loading control for comparative analysis.

### RNA-seq and analysis

For transcriptomic profiling, RPE-1 cells were cultured in 60 mm cell culture dishes (Thermo Scientific, 150462) under osmotic stress conditions for 6, 12, 24, 48 and 72 hr. At each time point, cells were harvested into a 1.5mL microcentrifuge tube after trypsinization, and immediately flash-frozen in liquid nitrogen before storage at −80°C until further processing. Total RNA was isolated from all samples in one sitting (all time points from three biological replicates) using the RNeasy Mini Kit (Qiagen, 74104) following the manufacturer’s protocol. RNA quantity and quality were preliminarily assessed with a Nanodrop 2000 Spectrophotometer (Thermo Scientific), before submission for sequencing. Library preparation and sequencing were performed by BGI Genomics (Hong Kong) on the BGISEQ platform with paired-end 100bp sequencing strategy. BGI Genomics’ Dr. Tom system was used for primary data processing. Low-quality reads, sequences containing adapters, and reads with high levels of N bases were filtered out from the raw data. Clean reads were aligned to the human reference genome (NCBI GCF_000001405.38_GRCh38.p12) using HISAT, and Bowtie2 was employed for transcript quantification. Differential expression analysis was performed using the R package DESeq2^30^. A time-series design incorporating condition, time, and the interaction between condition and time (∼ condition + time + condition:time) was used. Genes were pre-filtered to keep only those with at least 10 reads in at least three samples. To identify genes showing significant differences in temporal expression patterns between Hypo and Iso conditions, a Likelihood Ratio Test (LRT) was performed against a reduced model lacking the interaction term (∼ condition + time). Genes with an adjusted P-value < 0.05 were considered significant temporal genes. For visualization and clustering, raw counts were normalized using the Variance Stabilizing Transformation (VST). Principal Component Analysis (PCA) was performed on the VST data. K-means clustering (k = 6) was performed on the significant temporal genes using Z-scores calculated from the VST-normalized counts. Gene Ontology (GO) Biological Process enrichment analysis was conducted on each cluster using the compareCluster function in the R package clusterProfiler, utilizing the org.Hs.eg.db annotation database. GO results were simplified to reduce redundancy. Transcription factor (TF) activities were inferred using the R package decoupleR with the DoRothEA human regulon database (confidence levels A, B, and C)^33^. DESeq2 Wald statistics (t-values) for the Hypo vs Iso contrast at each time point were extracted and used as input for decoupleR. TF activities were estimated using the Univariate Linear Model (ULM) method. The top 40 TFs, selected based on the minimum P-value across all time points, were visualized.

### Chromosome spread

After 72 hr of osmotic treatments, p53^-/-^ and p21^-/-^ RPE-1 cells were arrested in mitosis with KaryoMAX Colcemid Solution (Gibco, 15210040, 1:100) for 4 hr. Cells were harvested by trypsinization, centrifuged at 1000rpm for 5 min, and resuspended in 1mL of culture medium. 5mL of 37°C pre-warmed 0.56-0.74% (w/v) potassium chloride was added dropwise, and the cell suspension was incubated at RT for 7-10 min. Cells were fixed with freshly prepared fixative, comprising 3:1 methanol:glacial acetic acid, with initial addition of 100µL per 10mL of cell suspension. After centrifugation at 1000rpm for 5 min, supernatant was removed, and cells were resuspended in 5-10mL of fresh fixative. Cells were fixed overnight at 4°C. Overnight fixed cells were warmed to RT for 2-3 min before centrifugation at 1000rpm for 5 min. Cells were further incubated with 5mL of fresh fixative at RT for 5-7 min before centrifugation and finally resuspension in 100-200µL of fixative. Pre-clean glass slides were steamed over boiling water for 5-10 sec to enhance droplet spreading. About 15µL of fixed cell suspension was dropped onto each slide from a height using a micropipette mounted on a retort stand. Two drops were dispensed per slide. Droplet boundaries were marked with a pencil. Spread quality was checked by a 10× phase-contrast microscope. Acceptable preparations were air-dried overnight at RT, before mounting with glass coverslips and Vectashield antifade mounting medium with DAPI (Vector Laboratories, H-1200). Samples were imaged on a Zeiss LSM 980 AiryScan confocal microscope with a 63× 1.4 N.A. oil immersion objective lens. Chromosomes were counted manually in a blinded manner in ImageJ. For each experimental group, around 30-40 well-spread metaphase prepared across three independent biological replicates were analyzed. Reliable chromosome spread could not be performed in WT RPE-1 cells because most cells were arrested in Hypo, which precluded observation of mitotic chromosomes in sufficient number.

### p53 synthesis and degradation

RPE-1 p53-mNG cells co-expressing H2B-mScarlet were generated and used in these experiments. Cells were treated with Iso or Hypo for 72 hr before seeding onto micropatterns for 2 hr before live-cell imaging. Imaging was performed using a Nikon W1 Spinning Disk system (Yokogawa CSU-W1) equipped with a 100× 1.4 N.A. oil immersion objective lens and a humidified chamber maintained at 37°C with 5% CO₂. Z-stack Images (10 µm range, 0.5 µm steps) were acquired every 10 min for 8-10 hr with the MetaMorph software. To measure the rate of p53 synthesis, cells were initially imaged for 30 min to establish baseline p53-mNG levels before adding 5 µM Nutlin, and continuing imaging for 8-10 hr. To monitor the rate of p53 degradation, seeded cells were pre-treated with 5 µM Nutlin for 24 hr. Nutlin-treated cells were imaged for 30 min to acquire baseline, followed by exchanging the Nutlin-containing media with non-drug containing media at the same osmotic stress, and continued imaging for 8-10 hr. Image analyses were performed with ImageJ with the TrackMate plugin for single-cell tracking. H2B-mScarlet intensity was used to generate a mask to measure nuclear p53mNG florescence intensities overtime. Three independent experiments were performed, with more than 10 cells analyzed per condition in each experiment.

### p53 nuclear transport

RPE-1 p53-mNG cells co-expressing H2B-mScarlet were prepared the same way as described earlier with the p53 synthesis and degradation experiments. Imaging and Fluorescence Loss in Photobleaching (FLIP) were performed with a Nikon W1 Laser Ablation Spinning Disk system (Yokogawa CSU-W1) with a 100× 1.4 N.A. oil immersion objective lens, housed in a humidified chamber maintained at 37°C with 5% CO₂. The scanning region of the 355nm photobleaching laser was pre-calibrated prior to the experiment. Regions of interest (ROI = 10000 pixel^2^) in the cytoplasm was defined, avoiding touching the nucleus. Cells were imaged for 1 min to acquire baseline, before FLIP was performed at the ROI by repeated cycles at 50% laser power and 10 iterations before image acquisition every 30 sec for 20 min. Images were acquired with the MetaMorph software and analyzed with ImageJ. Measurement for the rate of p53 nuclear import was attempted by fluorescence recovery after photobleaching (FRAP). In this experiment, the nuclei were either partially or totally bleached and the recovery of p53-mNG signal was monitored over time (data not shown). While recovery to a partially bleached region in the nuclei was quick due to rapid diffusion from existing nuclear pool, no appreciable increase in total p53-mNG nuclear signal was detected after 2 hr. This was likely due to the low abundance of cytoplasmic p53-mNG that was insufficient to replenish the loss of nuclear p53-mNG via nuclear import. Recovery of nuclear p53-mNG could only be achieved by new p53-mNG synthesis, which was beyond the FRAP observation window. The equilibrium state of p53 nuclear-cytoplasmic partition was instead estimated by quantifying the nuclear-cytoplasmic ratio of p53-mNG from fixed confocal images.

### FLIM-FRET measurements

Chromatin compaction in living cells were assayed via Fluorescence Lifetime Imaging - Förster Resonance Energy Transfer (FLIM-FRET) as previously described^43^. RPE-1 cells co-expressing H2B-mNeonGreen and H2B-mCherry were seeded onto 35 mm glass-bottom dishes (IWAKI, 3970-035) and treated with osmotic stress for 72 hr. FLIM was measured on a ZEISS LSM980 microscope equipped with a PicoQuant FLIM upgrade module and SymPhoTime64 software. Images were acquired with a 63× 1.4 N.A. oil immersion objective lens. Cells were maintained in a humidified chamber at 37°C and 5% CO_2_ during measurement. Data were collected for 60 sec per cell to ensure robust photon count. A control of only H2B-mNeonGreen expressing cells were included to establish the baseline fluorescence lifetime. A positive control of 100ng/mL TSA was included to induce chromatin decondensation, yielding the expected reduction in FRET efficiency and higher lifetime. At least 10 cells were captured per treatment condition, and the experiments were conducted in three biological replicates.

### Mathematical model

To quantitatively test the hypothesis that nuclear size reduction is sufficient to drive p53 activation, we developed a deterministic ordinary differential equation (ODE) model to describe the dynamics of the total nuclear p53RE reporter, 𝑅_𝑡𝑜𝑡_(𝑡). We assumed rapid nucleocytoplasmic transport equilibrium for total p53, 𝑃_𝑡𝑜𝑡_(𝑡), and the p53RE reporter, which allowed the use of a single-compartment varying-volume model. p53RE synthesis was assumed to be driven by nuclear p53 concentration, as defined by 𝑃_𝑡𝑜𝑡_(𝑡)/𝑉(𝑡), where 𝑉(𝑡) is the nuclear volume estimated from projected surface area measurements by raising them to the power of 3/2. Although the full system contains two coupled ODEs, the limited observed change in total nuclear p53 under both stress conditions allowed us to derive a reduced model by approximating the total p53 level as its time-averaged mean. This approximation enabled us to lump intrinsic synthesis and p53 levels into a single effective rate constant, 𝑘_𝑠𝑦𝑛,𝑒𝑓𝑓_. The dynamics of the normalized reporter, 𝑅̄(𝑡), is then described by the following simple ODE:

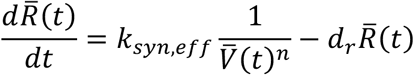

where 𝑉̄ (𝑡) is the normalized nuclear volume and 𝑑_𝑟_ is the degradation rate. We introduced a volume scaling exponent, *n*, to account for potential additional volume dependencies that may arise in confined biomolecular reactions. To determine the mechanism that best described our experimental observations, we defined four nested model classes (M0–M3) varying in complexity regarding *n* and condition-specific synthesis rates. We utilized Profile Likelihood Analysis to ensure parameter identifiability and an optimized data time-window analysis to select model M0 (where n=1), which explains the experimental data best while being identifiable and requiring the fewest free parameters. Please see the Supplementary Information for a more detailed description of the model, underlying equations and assumptions, and other statistical methods used for model selection.

### Statistical analysis

Statistical analyzes were performed using GraphPad Prism: t-tests for comparison between two groups, and one-way or two-way ANOVA followed by Tukey’s post hoc test for multiple comparisons. Unless otherwise stated, all data were represented as the average of at least three independent biological replicates, and the error bars were represented as the standard error of mean (SEM). ns represents insignificant difference where p > 0.05, * represents p ≤ 0.05, ** represents p ≤ 0.01, *** represents p ≤ 0.001 and **** represents p ≤ 0.0001.

## Data Availability

All raw RNA-seq data, as well as processed datasets, are available at the Gene Expression Omnibus database (GSEXXXXXX, to be released upon publication). All data generated or analyzed during this study are included in this article and its supporting supplementary files. Source data are provided with this paper. All other data are available from the corresponding authors upon reasonable request.

## Acknowledgement

We thank the Mechanobiology Institute of Singapore, specifically the Wet Lab Core for facility and equipment support, the Singapore Microscopy and Bioimage Analysis Core for microscopy support, the Nano and Microfabrication Core for fabricating the micropatterns, and the High Throughput Molecular Genetic Core for providing RNA-seq and flow cytometry support. We thank Teresa Ho and David Lane (A*STAR) for providing the RPE-1 p53-mNG cell line. We thank Paul Wan Sia Heng (NUS) for providing access to the osmometer for osmolarity measurements. We are grateful to Tetsuya Hiraiwa (Institute of Physics, Academia Sinica), Jacques Prost, Christophe Lamaze, Pierre Sens and Matthieu Piel (Institut Curie) for fruitful discussions and helpful inputs. This work was supported by the Singapore National Research Foundation’s Mid-Sized Grant (NRF-MSG-2023-0001).

## Author Contributions

B.S.W. and R.L. conceived the project and planned the experiments. B.S.W. performed most of the experiments. A.K.G. and S.X.S developed the mathematical model for the nuclear concentration-driven mechanism of p53 activation. H.C. prepared the micropattern and performed experiments related to p53 protein dynamics. K.L. and K.S. helped in performing confocal and live imaging experiments. B.A.J. developed the p53 activity reporter. Y.L. and J.Z. performed and analyzed the RNA-seq experiment. B.S.W. and A.K.G. analyzed data and prepared the figures. B.S.W. and R.L. wrote the manuscript with input from all co-authors.

---

**Supplementary Figure 1:**
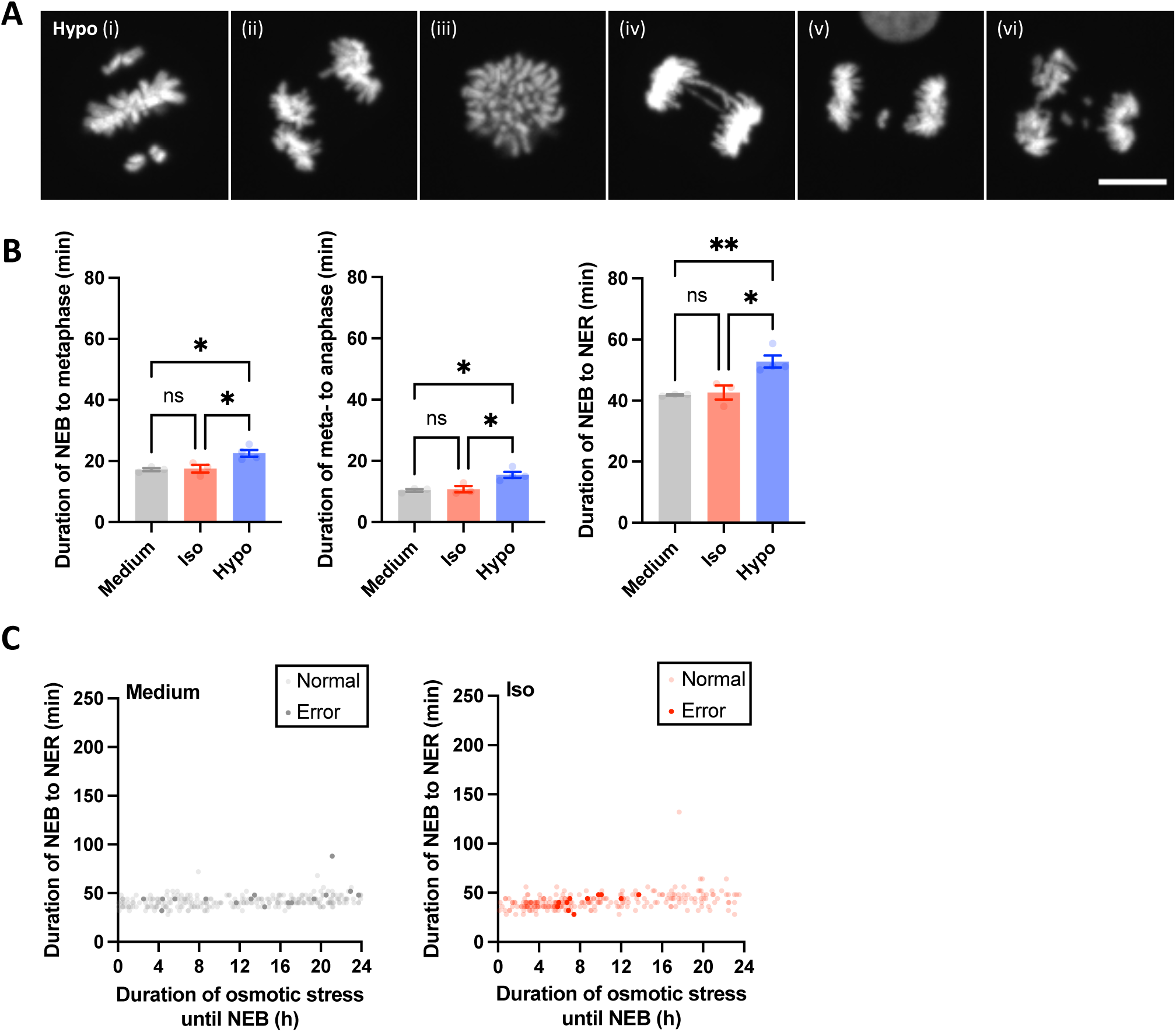
**(A)** Representative images of mitotic errors under Hypo condition: (i) unaligned chromosomes, (ii) multipolar mitosis, (iii) mitotic slippage, (iv) AB, (v) LC, and (vii) multiple defects (multipolar mitosis with LC). Scale bar = 10 µm. **(B)** Quantifications of the duration of NEB to metaphase (left), metaphase to anaphase (middle), and NEB to NER (right) under different osmotic conditions. **(C)** Correlation between the duration of NEB to NER (i.e. mitotic duration) and the duration osmotic stress exposure until NEB for all mitotic events under Medium (left) or Iso (right) conditions across all experimental replicates (n ≥ 3). Normal (light) and erroneous (dark) mitoses are differentially labeled. Bar graphs in (B) represent mean ± SEM from n ≥ 3 independent experiments. Data from each independent experiment were averaged to generate a single biological replicate value. Statistical significance was assessed via one-way ANOVA for (B). ns represents not significant, * represents p ≤ 0.05, ** represents p ≤ 0.01, *** represents p ≤ 0.001 and **** represents p ≤ 0.0001.

**Supplementary Figure 2:**
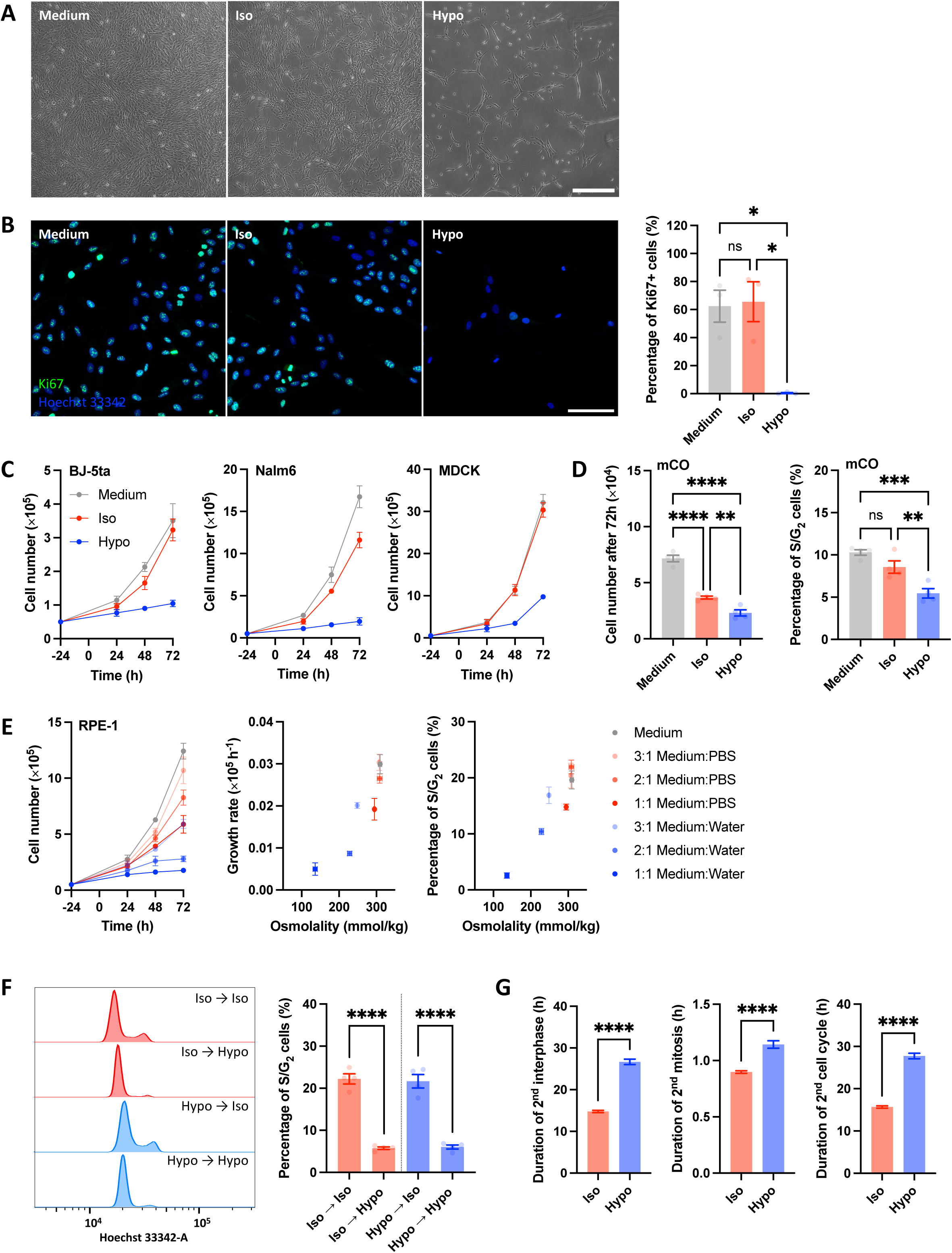
**(A)** Bright-field images of RPE-1 cells after 72 hr of osmotic stress. Scale bar = 500 µm. **(B)** Immunofluorescence images of RPE-1 cells stained with Ki67 antibody (green) and Hoechst 33342 (blue) after 72 hr of osmotic stress. Scale bar = 100 µm (left). Quantification of the percentage of Ki67+ cells (right). **(C)** Growth curves of BJ-5ta fibroblasts (left), MDCK epithelial cells (middle), and Nalm6 suspension cells (right) under different osmotic conditions. The dots and error bars represent mean ± SEM from n ≥ 3 independent experiments. **(D)** Quantifications of the cell number of mouse colonoid (mCO, left) and the percentage of S/G_2_ cells (right) after 72 hr of osmotic stress. **(E)** Growth curves of RPE-1 cells after 72 hr of treatment in media with varying osmolality (150-300 mmol/kg, left). Correlations between media osmolality and growth rates (middle) or the percentage S/G_2_ cells (right). The dots and error bars represent mean ± SEM from n ≥ 3 independent experiments. **(F)** Cell cycle profiles by genome content labeling 72 hr after switching to different osmotic conditions (left). RPE-1 cells were preconditioned in Iso or Hypo for 72 hr before the switch (i.e. Iso ◊ Hypo, Hypo ◊ Iso). Controls groups maintained in the same conditions were included (i.e. Iso ◊ Iso, Hypo ◊ Hypo). Quantification of the percentage of S/G_2_ cells 72 hr after the switch (right). **(G)** Quantifications of the duration of 2^nd^ interphase (left), 2^nd^ mitosis (middle), and 2^nd^ cell cycle (right) under Iso or Hypo conditions. Bar graphs in (B), (D), (F) and (G) represents mean ± SEM from n ≥ 3 independent experiments. For (B), data obtained from multiple fields of view from each independent experiment were averaged to generate a single biological replicate value. Statistical significance was assessed via one-way ANOVA for (B) and (D), and t-tests for (F) between the same precondition and (G). ns represents not significant, * represents p ≤ 0.05 and ** represents p ≤ 0.01, *** represents p ≤ 0.001 and **** represents p ≤ 0.0001.

**Supplementary Figure 3:**
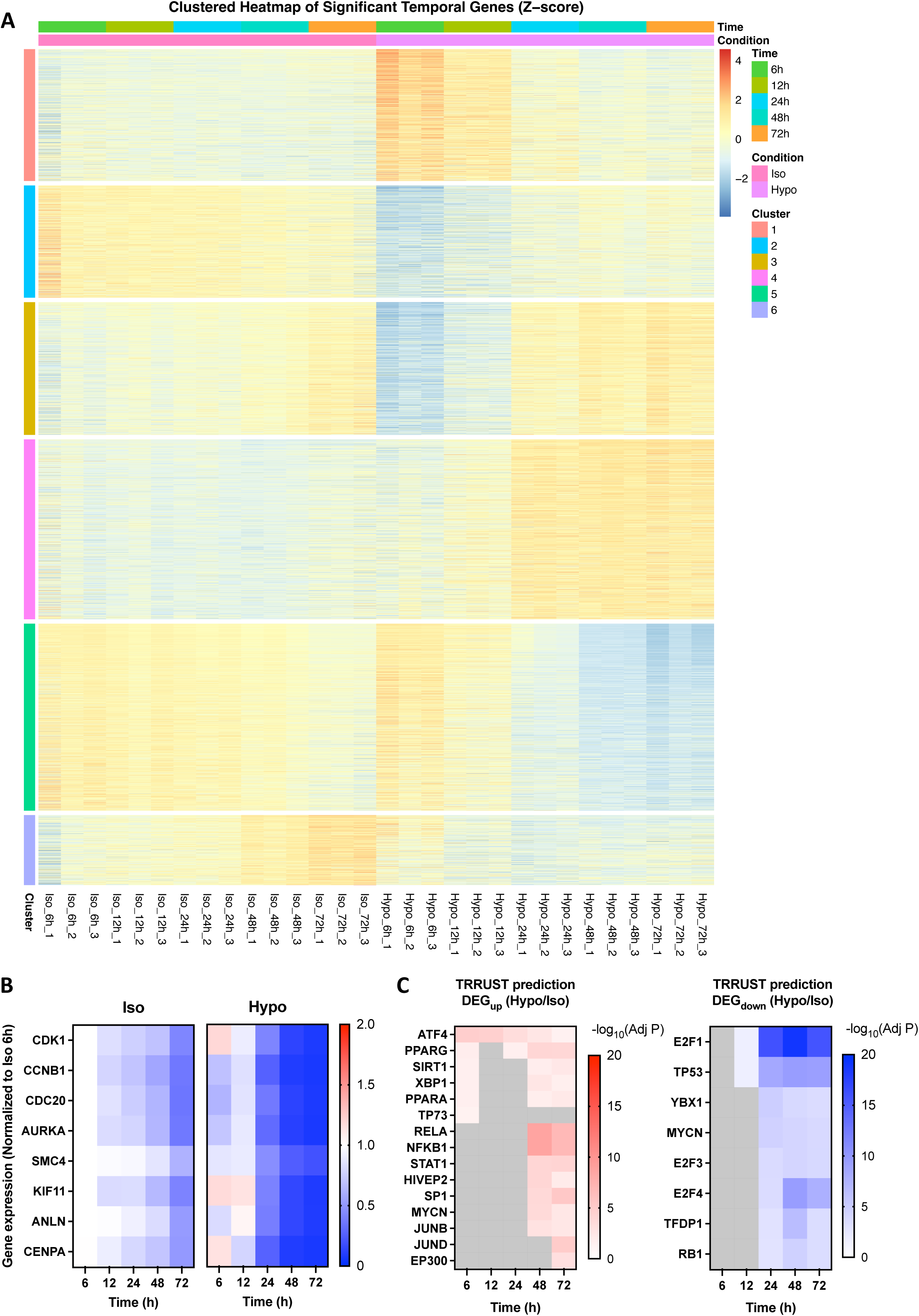
**(A)** Heatmap of significant temporal genes (identified via DESeq2 LRT) showing Z-score normalized expression (VST values). Genes (rows) are ordered according to the six clusters identified by K-means clustering (Cluster 1 to 6). **(B)** Heatmap of gene expression for selected genes associated with cell cycle and mitosis (i.e. *CDK1*, *CCNB1*, *CDC20*, *AURKA*, *SMC4*, *KIF11*, *ANLN* and *CENPA*) over time (6-72h) under Iso (left) or Hypo (right) conditions. Transcripts per million (TPM) values were normalized to Iso 6 hr (i.e. 1) for each gene and represented as the means of triplicates. **(C)** TRRUST transcription factor analyses of upregulated (DEG_up_, left) and downregulated differentially expressed genes (DEG_down_, right) over time (6-72h) between Hypo and Iso-treated samples (Hypo/Iso). Top predicted regulatory factors are shown. The heatmap represents −log_10_ (adjusted P value).

**Supplementary Figure 4:**
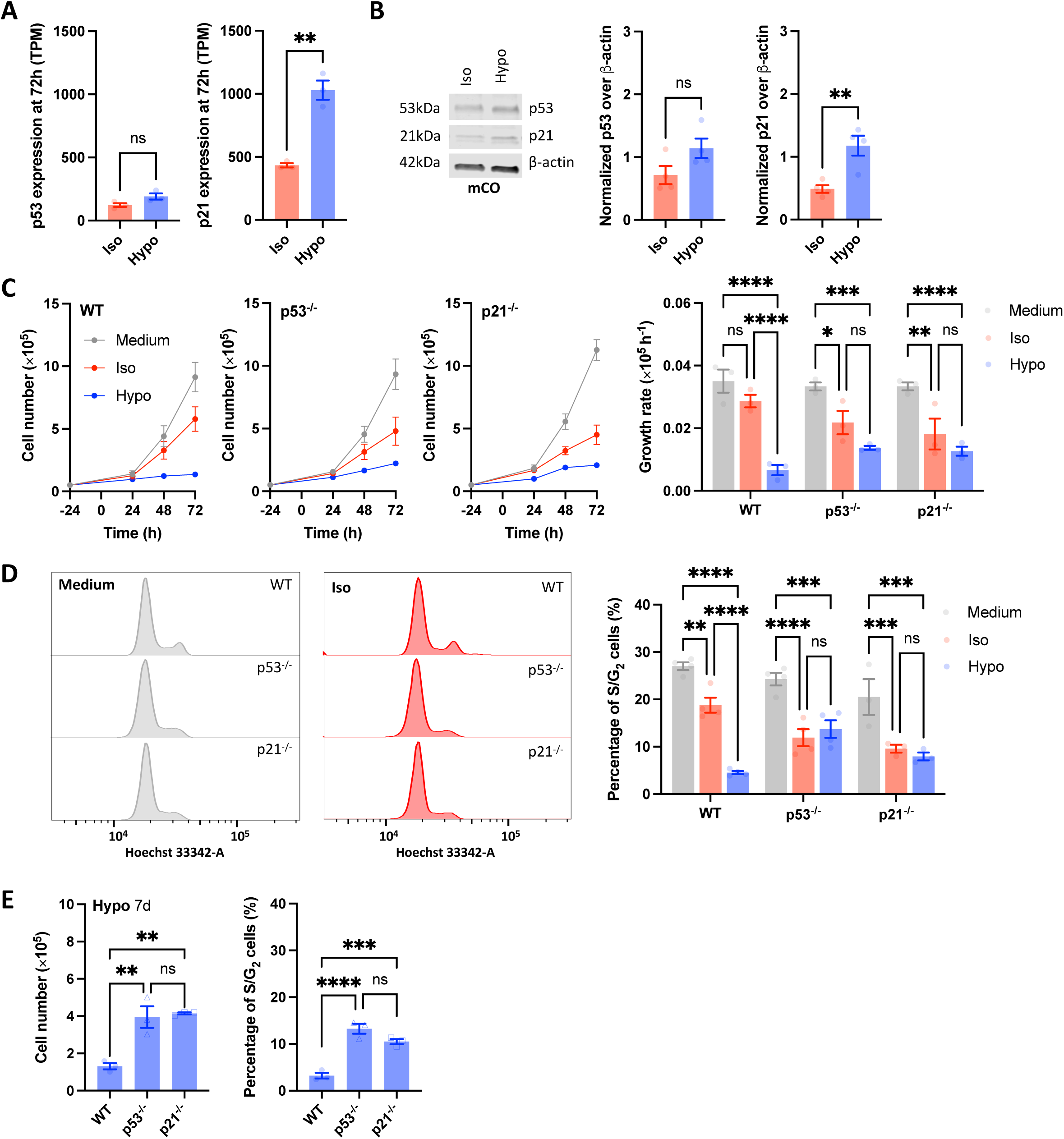
**(A)** mRNA expression levels (quantified as TPM from RNA-seq) of p53 (left) and p21 (right) after 72 hr of Iso or Hypo treatment. **(B)** Western blots of p53 and p21 in mCO after 72 hr of Iso or Hypo treatment (left). Quantifications of the protein level of p53 (middle) and p21 (left) normalized to β-actin. **(C)** Growth curves of WT, p53^-/-^ and p21^-/-^ cells in different osmotic conditions (left). The dots and error bars represent mean ± SEM from n ≥ 3 independent experiments. Growth rate comparisons were performed between different osmotic groups within each cell line (right). **(D)** Cell cycle profiles by genome content labeling after 72 hr of Medium (left) or Iso (middle) treatment in WT, p53^-/-^ and p21^-/-^ cells. Quantification of the percentage of S/G_2_ cells (right). Statistical comparisons were performed between different osmotic groups within each cell line. **(E)** Quantifications of the cell number (left) and the percentage of S/G_2_ cells (right) of WT, p53^-/-^ and p21^-/-^ cells after 7 days of Hypo treatment. All bar graphs represent mean ± SEM from n ≥ 3 independent experiments. Statistical significance was assessed via t-tests for (A) and (B), two-way ANOVA for (C) and (D), and one-way ANOVA for (E). ns represents not significant, * represents p ≤ 0.05, ** represents p ≤ 0.01, *** represents p ≤ 0.001 and **** represents p ≤ 0.0001.

**Supplementary Figure 5:**
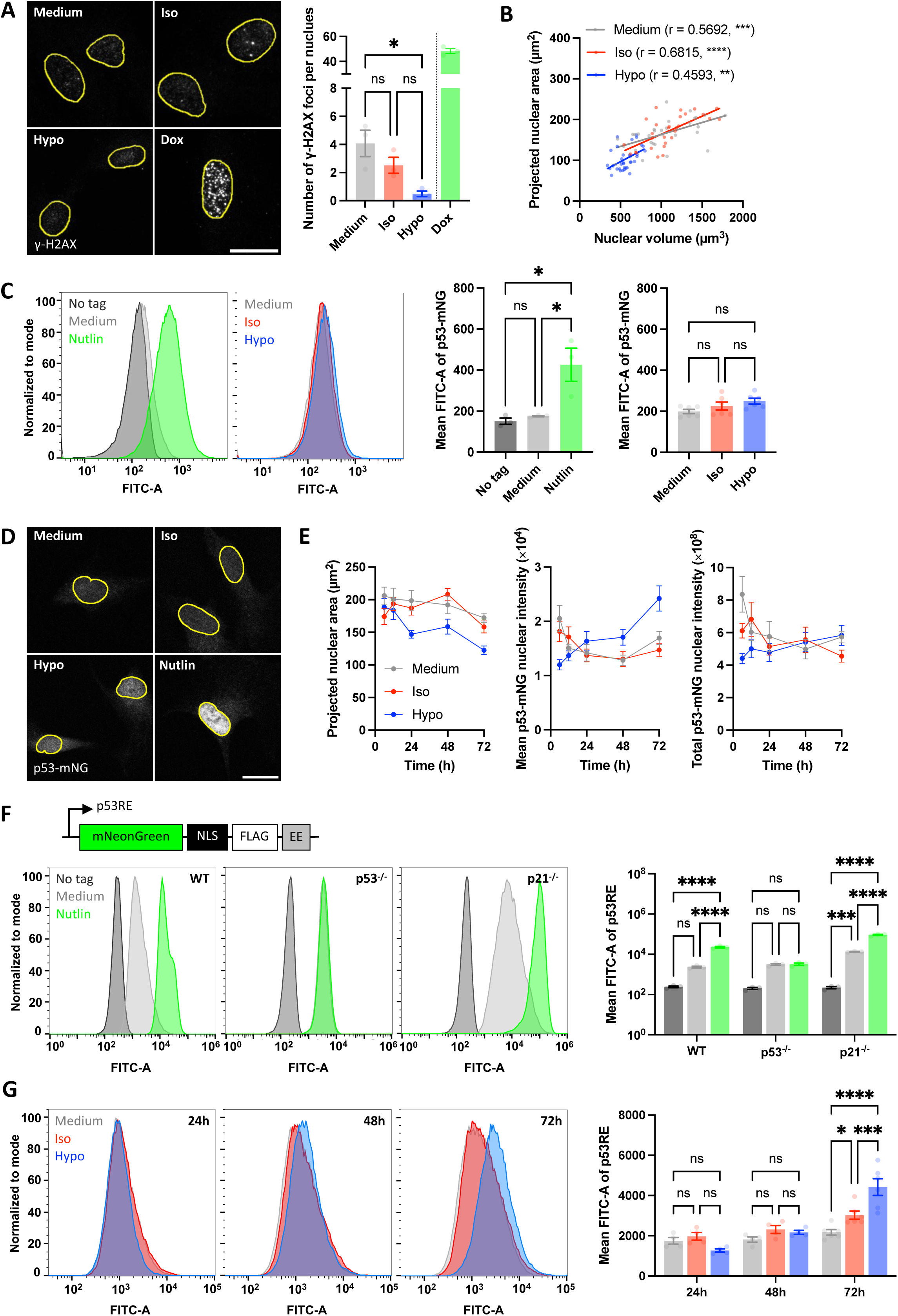
**(A)** Immunofluorescence confocal images of γ-H2AX in RPE-1 cells after 72 hr of osmotic stress. Treatment with 1 µM Doxorubicin (Dox) for 24 hr served as a DNA-damage positive control. Nuclei were counterstained with Hoechst 33342 (not shown) and outlined in yellow. Scale bar = 20 µm (left). Quantification of the number of γ-H2AX foci per nucleus (right). **(B)** Correlation between two-dimensional projected nuclear area and three-dimensional nuclear volume, measured from the same confocal images of DAPI-stained RPE-1 cells after 72 hr of osmotic stress. Solid lines represent linear regression fits. Pearson correlation coefficients (r) and their corresponding p-values are shown. **(C)** Flow cytometry histograms of RPE-1 cells expressing p53-mNG after 24 hr of 5 µM Nutlin treatment. WT cells were included as the no tag control, while RPE-1 p53-mNG cells in drug-free medium served as the unstimulated control (left). Flow cytometry histograms of RPE-1 p53-mNG cells after 72 hr of osmotic stress (middle left). Quantifications of the mean p53-mNG intensity following different treatments (middle right and right). **(D)** Fixed confocal images of RPE-1 p53-mNG cells after 72 hr of osmotic stress or 24 hr of 5 µM Nutlin treatment. Nuclei were counterstained with Hoechst 33342 (not shown) and outlined in yellow. Images represent z-projections from summed slices with matched intensity scaling across conditions. Scale bar = 20 µm. **(E)** Quantifications of the projected nuclear area (left), and the mean (middle) and total (right) p53-mNG nuclear intensity of RPE-1 p53-mNG cells fixed after 6, 12, 24, 48 and 72 hr of osmotic stress. Data represents mean ± SEM from ≥ 10 cells for each time point. **(F)** Plasmid design of the p53 activity reporter (p53RE). It comprises a p53 binding response element derived from the p21 promoter driving the expression of mNeonGreen tagged with a nuclear localizing sequence (NLS), a FLAG tag and a C-terminal EE degron (top). Flow cytometry histograms of RPE-1 WT, p53^-/-^ and p21^-/-^ cells expressing p53RE after 24 hr of 5 µM Nutlin treatment (bottom left), and quantification of mean p53RE intensity (bottom right). Untransduced cells were included as no tag controls, while p53RE-expressing cells in drug-free medium served as unstimulated controls. **(G)** Flow cytometry histograms of RPE-1 cells expressing p53RE after 24, 48 and 72 hr of osmotic stress (left), and quantification of mean p53RE intensity (right). All bar graphs represent mean ± SEM from n ≥ 3 independent experiments. For (A), data of multiple cells from each independent experiment were averaged to generate a single biological replicate value. Statistical comparisons for (A) were assessed between different osmotic conditions (excluding the positive control) with one-way ANOVA, via one-way ANOVA for (C), and via two-way ANOVA for (F) and (G). ns represents not significant, * represents p ≤ 0.05, ** represents p ≤ 0.01, *** represents p ≤ 0.001 and **** represents p ≤ 0.0001.

**Supplementary Figure 6:**
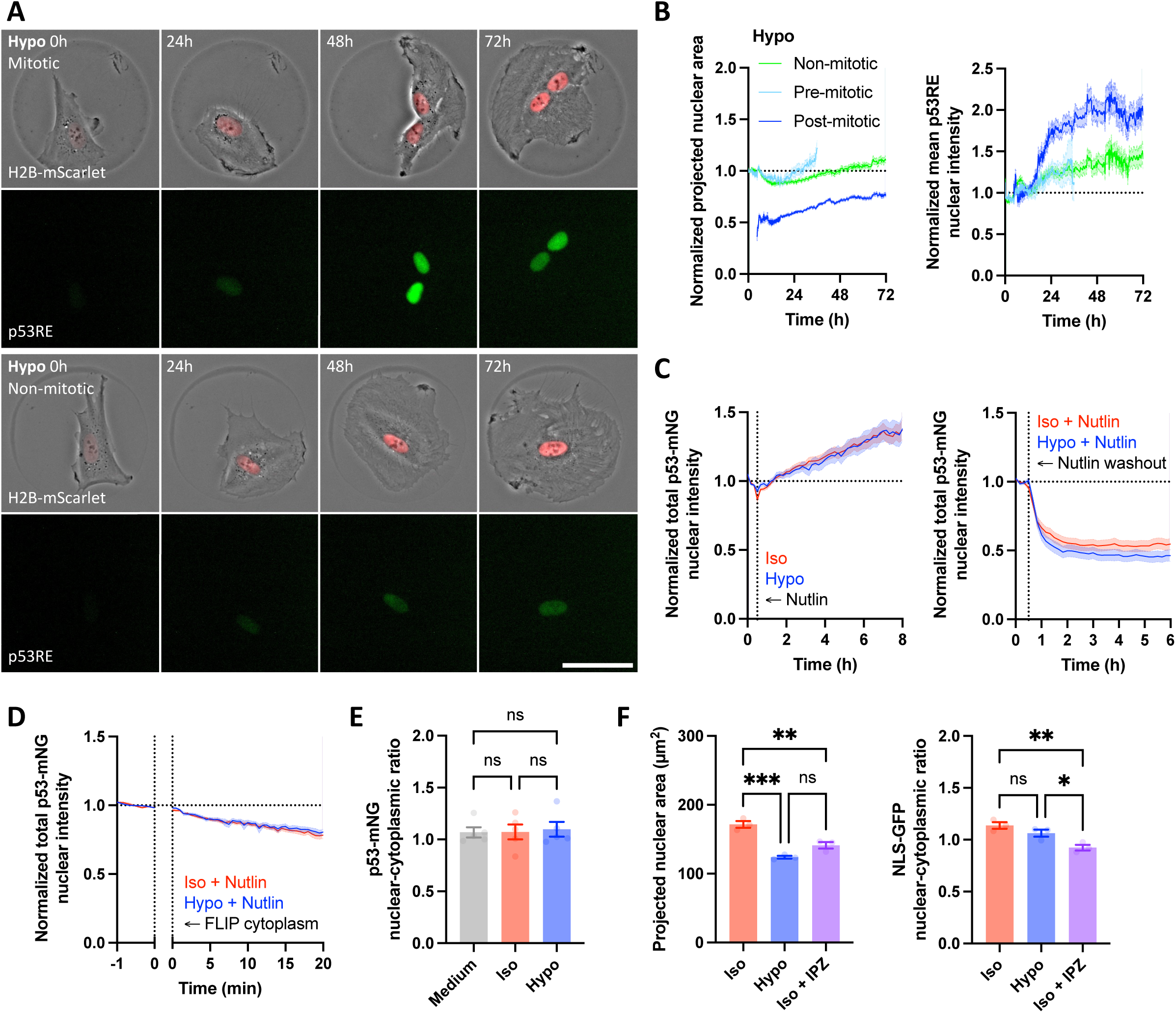
**(A)** Time-lapse montages of mitotic (top) and non-mitotic (bottom) RPE-1 cells co-expressing H2B-mScarlet and p53RE on fibronectin-coated micropatterns under Hypo condition for 72 hr. The montages were aligned to the start of imaging at 0 hr. Scale bar = 50 µm. **(B)** Quantifications of normalized projected nuclear area (left), and normalized mean p53RE nuclear intensity (right) over time. The data were normalized to the initial levels for each cell (average within 30 min from the start of imaging), and aligned to the start of imaging at 0 hr. A total of 67 pre-mitotic cells divided to form 134 post-mitotic daughter cells, and 60 non-mitotic cells were analyzed across four independent Hypo experiments. Line represents mean while shading represents SEM. **(C)** Quantification of the normalized total p53-mNG nuclear intensity of 72h-primed Iso or Hypo RPE-1 cells expressing p53-mNG before and after the addition of 5 µM Nutlin (left). Quantification of the normalized total p53-mNG nuclear intensity of 72h-primed Iso or Hypo RPE-1 cells expressing p53-mNG treated with 5 µM Nutlin for 24 hr before and after Nutlin washout (right). Line represents mean while shading represents SEM from n ≥ 3 independent experiments with at least 10 cells per replicate. **(D)** Quantification of the normalized total p53-mNG nuclear intensity of 72h-primed Iso or Hypo RPE-1 cells expressing p53-mNG treated with 5 µM Nutlin for 24 hr before and during Fluorescence Loss in Photobleaching. Line represents mean while shading represents SEM from n ≥ 3 independent experiments with at least 10 cells per replicate. **(E)** Quantification of the nuclear-to-cytoplasmic ratio of p53-mNG from confocal images of RPE-1 p53-mNG cells after 72 hr of osmotic stress depicted in Supplementary Figure 5D. **(F)** Quantifications of the projected nuclear area (left) and the nuclear-to-cytoplasmic ratio of NLS-GFP (right) from confocal images of RPE-1 cells expressing NLS-GFP after 72 hr of Iso, Hypo or Iso + IPZ treatments.

**Supplementary Figure 7:**
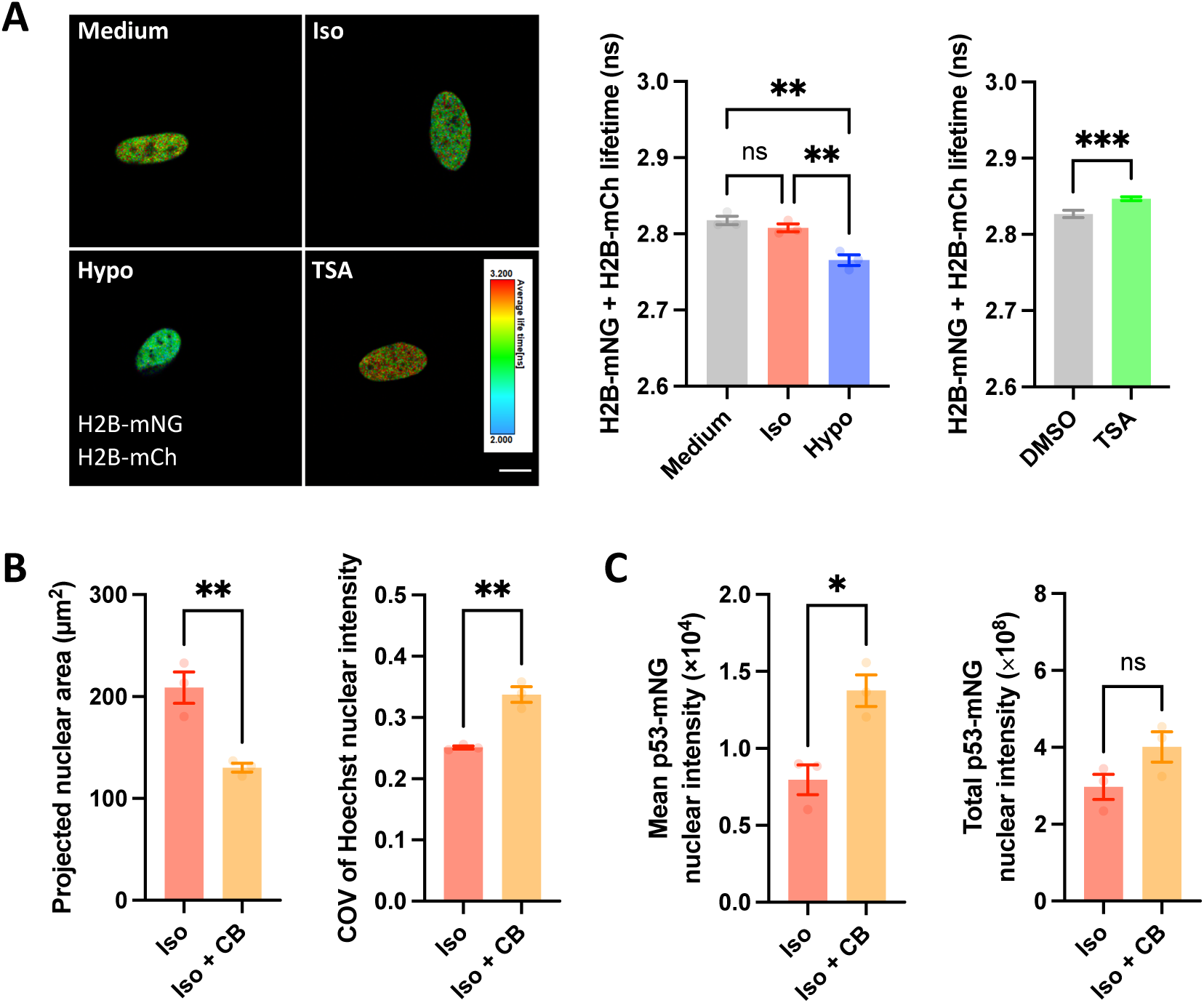
**(A)** Fluorescence Lifetime Imaging - Förster Resonance Energy Transfer (FLIM-FRET) images of RPE-1 cells expressing H2B-mNeonGreen (H2B-mNG) and H2B-mCherry (H2B-mCh) after 72 hr of osmotic treatment or 2 hr of 100ng/mL TSA (left). The lifetimes were represented by a heatmap with warmer colors indicating higher lifetime, lower FRET efficiency and less compact chromatin. Scale bar = 10 µm. Quantifications comparing the FRET-FLIM lifetime of Medium-, Iso- or Hypo-treated cells (middle), and DMSO- or TSA-treated cells (right). **(B)** Quantifications of the projected nuclear area (left) and the coefficient of variation (COV) of Hoechst nuclear intensity (right) after 72 hr of Iso treatment with or without 0.5µM CB-6644 (CB) in RPE-1 cells. **(D)** Quantifications of the mean (left) and total (right) p53-mNG nuclear intensity after 72 hr of Iso treatment with or without CB in RPE-1 cells expressing p53-mNG. All bar graphs represent mean ± SEM from n ≥ 3 independent experiments. Data of multiple cells from each independent experiment were averaged to generate a single biological replicate value. Statistical comparisons were assessed via one-way ANOVA for (A-middle), and via t-tests for (A-right), (B) and (C). ns represents not significant, * represents p ≤ 0.05, ** represents p ≤ 0.01, *** represents p ≤ 0.001 and **** represents p ≤ 0.0001.

**Movie 1:** Mitotic progression of RPE-1 cells expressing H2B-mNeonGreen under Medium, Iso, or Hypo conditions. Images were acquired every 4 min for 24 hr.

**Movie 2:** Cell cycle dynamics of FUCCI-expressing RPE-1 cells on fibronectin-coated circular micropatterns under Iso condition. Images were acquired every 10 min for 72 hr.

**Movie 3:** Cell cycle dynamics of FUCCI-expressing RPE-1 cells on fibronectin-coated circular micropatterns under Hypo condition. Images were acquired every 10 min for 72 hr.

**Movie 4:** p53 activity and nuclear dynamics of RPE-1 cells expressing p53RE and H2B-mScarlet on fibronectin-coated circular micropattern undergoing cell division under Iso condition. Images were acquired every 10 min for 72 hr.

**Movie 5:** p53 activity and nuclear dynamics of RPE-1 cells expressing p53RE and H2B-mScarlet on fibronectin-coated circular micropattern undergoing cell division under Hypo condition. Images were acquired every 10 min for 72 hr.

**Movie 6:** p53 activity and nuclear dynamics of RPE-1 cells expressing p53RE and H2B-mScarlet on fibronectin-coated circular micropattern undergoing cell division under Iso condition with 10 µM Importazole (IPZ). Images were acquired every 10 min for 72 hr.

**Movie 7:** p53 activity and nuclear dynamics of RPE-1 cells expressing p53RE and H2B-mScarlet on fibronectin-coated circular micropattern undergoing cell division under Iso condition with 0.5µM CB-6644 (CB). Images were acquired every 10 min for 72 hr.

**Supplementary Movie 1:** p53 activity and nuclear dynamics of non-mitotic RPE-1 cells expressing p53RE and H2B-mScarlet on fibronectin-coated circular micropattern under Hypo condition. Images were acquired every 10 min for 72 hr.

