## Supplementary Information for "p53 preserves genome stability in dividing cells under hypo-osmotic stress"

### Supplementary Information (SI)

We aim to develop a simple mathematical model that describes the post-mitotic time-dependent dynamics of the p53 activity reporter (p53RE) for dividing cells under iso-osmotic (Iso) and hypo-osmotic (Hypo) stress.

#### General Framework and Normalization

We begin with a two-component model describing the dynamics of the total amount of nuclear p53 protein, $P_{tot}(t)$, and the total amount of p53RE, $R_{tot}(t)$. We assume a single-compartment model where both proteins primarily reside and function within the nucleus with volume $V(t)$ and rapid transport kinetics equilibrium that can be excluded from the model.

The general ODE describing the dynamics of $P$ and $R$ are as follows

$$\frac{dP_{tot}(t)}{dt} = k_p - d_p P_{tot}(t)$$

$$\frac{dR_{tot}(t)}{dt} = k_r \frac{P_{tot}(t)}{V(t)^n} - d_r R_{tot}(t)$$

The above equations describe the time rate of change of p53 and p53RE, which are represented by the differences in the rates of synthesis and degradation. The rate of synthesis of p53RE ($R_{tot}(t)$), is assumed to follow a simple mass-action kinetics with respect to p53 concentration in the nucleus, $P_{tot}(t)/V(t)$, and additional volume effect (captured by the exponent n) coming from nonideal conditions in the nucleus (this is explained in detail in the next section).

To facilitate comparison with experimental data, we define normalized quantities (denoted by an overhead bar) relative to their respective values at $t = 0$, the time of Nuclear Envelope Reformation (NER):

$$\bar{P}(t) = \frac{P_{tot}(t)}{P_{tot}(0)}; \;\; \bar{R}(t) = \frac{R_{tot}(t)}{R_{tot}(0)}; \;\; \bar{V}(t) = \frac{V(t)}{V(0)}$$

The general ODE describing the dynamics of $P$ and $R$ are as follows

$$\frac{d\bar{P}}{dt} = \alpha_p - d_p \bar{P}(t)$$

$$\frac{d\bar{R}}{dt} = k_{\text{syn},0} \frac{\bar{P}(t)}{\bar{V}(t)^n} - d_r \bar{R}(t)$$

with

$$\alpha_p = \frac{k_p}{P_{\text{tot}}(0)}, \qquad k_{\text{syn},0} = k_r \frac{P_{\text{tot}}(0)}{R_{\text{tot}}(0) V(0)^n}$$

Experimentally, within a chosen time window ($[t_1, t_2]$), ($\overline{P}(t)$) is observed to vary slowly and approximately linearly with a similar slope in Iso and Hypo conditions. We define the time-averaged normalized p53 over the analysis window:

$$\overline{P}_* \equiv \frac{1}{T}\int_{t_1}^{t_2} \overline{P}\,(t)dt, \qquad T = t_2 - t_1$$

For example, for an approximately linear drift ($\overline{P}(t) \approx 1 + s(t - t_1)$),

$$\overline{P}_* = 1 + \frac{sT}{2}$$

i.e. ($\overline{P}_*$) coincides with ($\overline{P}$) at the mid-time of the window.

The time-averaged p53 approximation consists of replacing ($\overline{P}(t)$) in the reporter equation by the constant ($\overline{P}_*$):

$$\frac{d\overline{R}_{\text{avg}}}{dt} = k_{\text{syn,eff}}\frac{1}{\overline{V}(t)^n} - d_r\overline{R}_{\text{avg}}(t)$$

with the effective synthesis parameter

$$k_{\text{syn,eff}} = k_{\text{syn,0}}\overline{P}_*$$

This yields a reduced one-equation model in which differences between conditions (e.g. Iso vs Hypo) enter through ($V(t)$), ($n$), and any condition-specific modulation of ($k_{\text{syn,eff}}$), while the measured slow drift of p53 is consistently absorbed into a single effective parameter. In the next section, we derive an analytic form of the correction term that is needed for this approximation and show that this is small under current experimental conditions.

Applying this assumption simplifies the system to a single ODE for the reporter dynamics. The collection of constant terms related to initial protein levels and the intrinsic reporter synthesis rate ($k_r$) is consolidated into a single, effective synthesis rate constant, $k_{syn,eff}$:

$$k_{syn,eff} = k_r\frac{\overline{P}_*}{R_{tot}(0)V(0)^n}$$

To generalize this for hypothesis testing, we introduce a volume scaling exponent, $n$. This exponent allows us to test whether the synthesis rate is driven purely by concentration ($n = 1$) or by a more complex, non-linear function of volume. This yields the final, generalized model used for fitting:

$$\frac{d\bar{R}(t)}{dt} = k_{syn,eff}\frac{1}{\overline{V}(t)^n} - d_r\bar{R}(t)$$

Where:

- $\overline{R}(t)$: Normalized total p53RE reporter.

- $\overline{V}(t)$: Normalized nuclear volume.
- $k_{syn}$: The lumped, normalized synthesis rate constant.
- $n$: The volume scaling exponent (a key parameter for hypothesis testing).
- $d_r$: The first-order degradation rate constant

This single ODE forms the basis of the hierarchical model comparison.

**Volume Dependence of Reaction Rates**

Standard mass action kinetics are typically formulated under the implicit assumption of a constant reaction volume. However, in our system, the nuclear volume changes significantly during the post-mitotic expansion phase and differs between Iso and Hypo conditions. To accurately model the rate of p53RE synthesis under these conditions, the reaction rate must be formulated to explicitly account for volume dependence.

Consider a simple elementary bimolecular reaction where species A and B react to form C:

$$A + B \overset{k}{\rightarrow} C$$

The reaction velocity $v$ (defined as concentration change per unit time in a static volume) is given by the law of mass action:

$$v = k[A][B]$$

where $k$ is the rate constant, and $[A]$ and $[B]$ are the molar concentrations of the reactants. When considering a system with time-dependent volume $V(t)$, it is physically more robust to track the total number of molecules, $N_C$. The rate of change of the number of product molecules (the extensive reaction rate) is the reaction velocity integrated over the volume:

$$\frac{dN_C}{dt} = v \cdot V(t) = k[A][B]V(t)$$

Substituting the definition of concentration, $[X] = N_X/V(t)$, where $N_X$ is the total number of molecules of species $X$, we obtain:

$$\frac{dN_C}{dt} = k\left(\frac{N_A}{V(t)}\right)\left(\frac{N_B}{V(t)}\right)V(t)$$

Simplifying this expression yields:

$$\frac{dN_C}{dt} = k\frac{N_A N_B}{V(t)}$$

Thus, for a diffusion-limited bimolecular reaction within a confined compartment, the probability of collision and the effective reaction rate scales inversely with the volume ($V^{-1}$).

**Application to p53 Transcriptional Activity**

We apply this framework to the synthesis of p53RE. The rate-limiting step is conceptualized as the binding of the transcription factor p53 ($P$) to its specific promoter sites on the DNA to initiate transcription. Here, $N_A$ corresponds to the total number of nuclear p53 molecules ($P_{tot}$) and $N_B$ corresponds to the number of accessible gene loci ($N_{gene}$), which is effectively constant (assuming diploidy and constant gene copy number during $G_1$).

The rate of reporter production ($R_{prod}$) can therefore be expressed as:

$$R_{prod} \propto \frac{P_{tot} \cdot N_{gene}}{V(t)}$$

This is assuming that the search time for the transcription factor to find the promoter is the rate-limiting step and is diffusion-controlled, the effective association rate scales inversely with the search volume.

Since $N_{gene}$ is constant, the rate scales as:

$$R_{prod} \propto \frac{P_{tot}}{V(t)}$$

This corresponds to the standard concentration-dependent synthesis assumed in Model M0 (where the volume exponent $n = 1$).

While ideal bimolecular collision theory predicts a strict inverse volume dependence ($V^{-1}$), the nuclear environment in a cell is more complex. Factors such as macromolecular crowding, fractal-like chromatin architecture, or the requirement for multi-protein complex assembly could cause effective reaction rates to deviate from simple mass action kinetics. To capture these potential non-linearities without over-specifying the biophysics, we generalized the synthesis term using an empirical volume scaling exponent, $n$:

$$\frac{dN_{p53RE}}{dt} = k_{syn} \frac{P_{tot}}{V(t)^n} - d_r N_{p53RE}$$

Here, $n = 1$ represents ideal concentration-driven kinetics, while deviations ($n \neq 1$) account for anomalous diffusion or higher-order reaction dependencies within the changing nuclear compartment. This formulation lumps the multi-step gene expression (promoter search and binding, initiation, elongation, translation) into a single synthesis rate. If a volume-independent step (e.g., polymerase elongation or mRNA export) were the primary bottleneck, the system would effectively show no dependence on volume ($n \approx 0$). However, our model selection analysis (described later) strongly favors $n \approx 1$, supporting the hypothesis that the concentration-dependent probability of p53 finding and binding its promoter is the rate-limiting step sensitive to osmotic stress in this context.

**Analytic Form of The Correction Term**

Here we derive the analytical form of the correction term arising when replacing total p53 by its time average. Let ($\overline{R}_{\text{exact}}(t)$) solve the full two-component driven equation with time-dependent ($\overline{P}(t)$):

$$\frac{d\overline{R}}{dt} + d_r \overline{R}(t) = k_{\text{syn,0}} \frac{\overline{P}(t)}{\overline{V}(t)^n}$$

For ($t \in [t_1, t_2]$), the exact solution is

$$\overline{R}_{\text{exact}}(t) = e^{-d_r(t-t_1)} \left[ \overline{R}(t_1) + k_{\text{syn,0}} \int_{t_1}^{t} e^{d_r(\tau - t_1)} \frac{\overline{P}(\tau)}{\overline{V}(\tau)^n} d\tau \right]$$

The time-averaged model replaces ($\overline{P}(\tau)$) by ($\overline{P}_*$):

$$\overline{R}_{\text{avg}}(t) = e^{-d_r(t-t_1)} \left[ \overline{R}(t_1) + k_{\text{syn,0}} \overline{P}_* \int_{t_1}^{t} e^{d_r(\tau - t_1)} \frac{1}{\overline{V}(\tau)^n} d\tau \right]$$

Their difference defines the perturbative correction term:

$$\Delta\overline{R}(t) \equiv \overline{R}_{\text{exact}}(t) - \overline{R}_{\text{avg}}(t) = e^{-d_r(t-t_1)} k_{\text{syn,0}} \int_{t_1}^{t} e^{d_r(\tau - t_1)} \frac{\overline{P}(\tau) - \overline{P}_*}{\overline{V}(\tau)^n} d\tau$$

Equivalently, writing ($\overline{P}(\tau) = \overline{P}_* + \delta P(\tau)$) with ($\int_{t_1}^{t_2} \delta\, P(\tau) d\tau = 0$),

$$\Delta\overline{R}(t) = e^{-d_r(t-t_1)} k_{\text{syn,0}} \int_{t_1}^{t} e^{d_r(\tau - t_1)} \frac{\delta P(\tau)}{\overline{V}(\tau)^n} d\tau$$

This expression is the analytic first-order correction to the reduced model arising from p53 non-constancy.

We now quantify when ($\Delta\overline{R}(t)$) is small compared to the reporter dynamics and define the maximal fractional deviation of p53 from its time average:

$$\delta_P \equiv \max_{t \in [t_1, t_2]} \left| \frac{\overline{P}(t) - \overline{P}_*}{\overline{P}_*} \right|$$

Let ($\overline{V}_{\text{min}}^n = \min_{[t_1, t_2]} \overline{V}(t)^n$) and let

$$\overline{R}_{\text{scale}} \sim \frac{k_{\text{syn,0}} \overline{P}_*}{d_r \overline{V}_{\text{typ}}^n}$$

denote the characteristic reporter amplitude in this window, with ($\overline{V}_{\text{typ}}$) a typical volume factor.

From the correction integral above,

$$|\Delta\overline{R}(t)| \leq k_{\text{syn,0}} \overline{P}_* \delta_P \frac{1 - e^{-d_r(t-t_1)}}{d_r \overline{V}_{\text{min}}^n}$$

Hence the fractional error satisfies

$$\frac{|\Delta\overline{R}(t)|}{\overline{R}_{\text{scale}}} \lesssim \delta_P \cdot \frac{\overline{V}_{\text{typ}}^n}{\overline{V}_{\text{min}}^n} \cdot (1 - e^{-d_r(t-t_1)}) \sim O(\delta_P)$$

i.e. the error is of the same order as the relative variation of p53 around its mean.

A complementary, dynamical criterion uses the drift rate of p53. Over one reporter lifetime ($1/d_r$), the normalized p53 changes by approximately

$$\Delta\overline{P}_{\text{life}} \approx \left|\frac{d\overline{P}}{dt}\right| \frac{1}{d_r}$$

so a sufficient condition for the time-averaged approximation to be accurate is

$$\frac{\Delta\overline{P}_{\text{life}}}{\overline{P}_*} = \frac{1}{\overline{P}_*} \left|\frac{d\overline{P}}{dt}\right| \frac{1}{d_r} \ll 1$$

The time-averaged p53 approximation (and resulting one-equation reporter model with ($k_{\text{syn,eff}} = k_{\text{syn,0}}\overline{P}_*$)) is a controlled and quantitatively accurate reduction of the full two-component system when:

1. Small relative drift over the analysis window

$$\max_{t\in[t_1,t_2]} \left|\frac{\overline{P}(t) - \overline{P}_*}{\overline{P}_*}\right| \ll 1$$

2. Slow drift compared to reporter turnover

$$\frac{1}{\overline{P}_*} \left|\frac{d\overline{P}}{dt}\right| \frac{1}{d_r} \ll 1$$

These are reasonable assumptions based on our experimental results of the rates of change of total p53RE and p53 protein over time (Figure 5F and Supplementary Figure 5E).

**Hierarchical Model Classes (M0-M3)**

To systematically test the "volume-only" hypothesis, we implemented a nested hierarchy of four model classes (M0, M1, M2, M3). These models introduce increasing complexity, allowing us to use statistical tests to justify the inclusion of additional parameters. All models are fitted simultaneously to data from two experimental conditions: Iso and Hypo.

**Model M0: Concentration-Only**

This is the simplest baseline model, which assumes that the synthesis rate is purely dependent on p53 concentration and that this relationship is identical across both Iso and Hypo conditions.

- Hypothesis: Volume changes are sufficient to explain the dynamics, with a direct 1:1 relationship between concentration and synthesis rate.
- ODE:

$$\frac{d\bar{R}(t)}{dt} = k_{syn,shared}\frac{1}{\bar{V}(t)} - d_r\bar{R}(t)$$

- Parameters:
  - $k_{syn,shared}$ $(h^{-1})$ Fitted, shared synthesis rate for both iso and hypo conditions.
  - $n$ fixed to 1.0
  - $d_r$ $(h^{-1})$ fixed

**Model M1: Shared Volume Exponent**

This model extends M0 by allowing the volume scaling exponent, n, to be a free parameter. This tests whether the relationship between volume and synthesis deviates from a simple concentration model, but assumes this relationship is the same for both conditions.

- Hypothesis: Volume has a consistent, non-linear effect on synthesis rate across conditions.
- ODE:

$$\frac{d\bar{R}(t)}{dt} = k_{syn,shared}\frac{1}{\bar{V}(t)^{n_{shared}}} - d_r\bar{R}(t)$$

- Parameters:
  - $k_{syn,shared}$ $(h^{-1})$ Fitted, shared synthesis rate for both iso and hypo conditions.
  - $n_{shared}$ Fitted, shared volume exponent.
  - $d_r$ $(h^{-1})$ fixed

**Model M2: Condition-Specific Amplitude**

This model tests whether the two conditions (Iso versus Hypo) have different intrinsic synthesis rates, while assuming a simple concentration-driven mechanism (n = 1).

- Hypothesis: The osmotic stress condition itself, independent of volume changes, alters the p53 transcriptional activity.
- ODEs:
  - Iso: $\frac{d\bar{R}_{iso}(t)}{dt} = k_{syn\_iso} \cdot \frac{1}{\bar{V}_{iso}(t)} - d_r \cdot \bar{R}_{iso}(t)$
  - Hypo: $\frac{d\bar{R}_{hypo}(t)}{dt} = k_{syn\_hypo} \cdot \frac{1}{\bar{V}_{hypo}(t)} - d_r \cdot \bar{R}_{hypo}(t)$
- Parameters:
  - $k_{syn,iso}$ $(h^{-1})$ Fitted, synthesis rate for Iso.
  - $k_{syn,hypo}$ $(h^{-1})$ Fitted, synthesis rate for Hypo.
  - $n$ fixed to 1.0
  - $d_r$ $(h^{-1})$ fixed

#### Model M3: Flexible Volume Model

This is the most complex model in the hierarchy, combining the features of M1 and M2. It allows for both a non-linear volume scaling effect and condition-specific synthesis rates.

- Hypothesis: Both a non-linear volume effect and a condition-specific transcriptional response are required to explain the data.
- ODEs:
  - Iso: $\frac{d\overline{R}_{iso}(t)}{dt} = k_{syn\_iso} \cdot \frac{1}{\overline{V}_{iso}(t)^{n_{shared}}} - d_r \cdot \overline{R}_{iso}(t)$
  - Hypo: $\frac{d\overline{R}_{hypo}(t)}{dt} = k_{syn\_hypo} \cdot \frac{1}{\overline{V}_{hypo}(t)^{n_{shared}}} - d_r \cdot \overline{R}_{hypo}(t)$
- Parameters:
  - $k_{syn,iso}$ ($h^{-1}$) Fitted, synthesis rate for Iso.
  - $k_{syn,hypo}$ ($h^{-1}$) Fitted, synthesis rate for Hypo.
  - $n_{shared}$ Fitted, shared volume exponent.
  - $d_r$ ($h^{-1}$) fixed

#### Model Fitness Criteria

The model is fitted to experimental data by minimizing a chi-squared ($\chi^2$) objective function. To ensure a balanced fit across both Iso and Hypo conditions, the optimizer minimizes the *maximum* of the two individual $\chi^2$ values ($\chi^2_{obj} = \max(\chi^2_{iso}, \chi^2_{hypo})$), subject to a penalty if either value is unacceptably high.

However, for model selection, the statistically appropriate measure is the *sum* of the chi-squared values from the independent datasets. This $\chi^2_{sum}$ is calculated for the best-fit parameter set and used to compute the Akaike Information Criterion (AIC) and Bayesian Information Criterion (BIC)[1].

- Chi-Squared Sum:

$$\chi^2_{sum} = \chi^2_{iso} + \chi^2_{hypo} = \sum_{i \in iso} \left( \frac{R_{pred}(t_i) - R_{obs}(t_i)}{SE(t_i)} \right)^2 + \sum_{j \in hypo} \left( \frac{R_{pred}(t_j) - R_{obs}(t_j)}{SE(t_j)} \right)^2$$

- AIC and BIC Calculation:

$$AIC = \chi^2_{sum} + 2k$$

$$BIC = \chi^2_{sum} + k \ln N$$

  - $k$: Number of fitted parameters in the model.
  - $N$: Total number of data points across both conditions in the fitting window.

AIC and BIC are used to rank the models. They provide a principled trade-off between goodness-of-fit and model complexity. A lower AIC/BIC score indicates a more parsimonious model that better explains the data without overfitting. The difference in AIC/BIC scores

($\Delta AIC$, $\Delta BIC$) between models is used to determine the level of support for one model over another.

For datasets with a limited number of data points relative to the number of fitted parameters, AIC can tend to favor overly complex models. The corrected Akaike Information Criterion (AICc) adjusts for this by applying a greater penalty for each additional parameter.

$$\text{AICc} = \text{AIC} + \frac{2k(k+1)}{N-k-1}$$

Substituting the expression for AIC, this becomes:

$$\text{AICc} = \chi^2_{\text{sum}} + 2k + \frac{2k(k+1)}{N-k-1}$$

**Nested Model Comparison and Selection**

Next, we performed a systematic comparison of the four nested model classes (M0-M3), to determine the most appropriate model to explain the observed p53RE dynamics. Our goal was to select the most parsimonious model that adequately describes the data, while ensuring the model's parameters are robust and interpretable.

We first assessed the degree to which each model could fit the experimental data. The simplest model, M0, assumes that p53RE synthesis is dependent only on p53 concentration (i.e., inversely proportional to nuclear volume, with a fixed exponent of 1). The more complex models add parameters to account for a variable volume exponent (M1), condition-specific synthesis rates (M2), or both (M3).

As shown by the representative fit for Model M0 (SI Figure 1A), even the simplest model provides an excellent fit for the experimental data for both Iso and Hypo conditions. The model correctly captures the relatively stable p53RE concentration in Iso and the increasing trend in Hypo. The more complex models – M1 (SI Figure 1B), M2 (SI Figure 1C) and M3 (SI Figure 1D) show better fit as indicated by their lower AIC values and higher $R^2$.

We also tested how the fitting for each of the models changes for different p53RE degradation rates (SI Figure 2A-B). We found the more complex models' (M2 and M3) AIC values are quite sensitive to this degradation rate while M0 is the least sensitive.

While all models fit the data well, information criteria like AIC are designed to penalize model complexity. The AIC vs. R-squared plot (SI Figure 2C) visualizes the trade-off between goodness-of-fit ($R^2$, higher is better) and the information criterion (AICc, lower is better).

This plot reveals two key insights:

- The more complex models (M1, M2, and M3) cluster in the bottom-right, indicating they are formally "better" models according to the AIC metric and achieve a slightly higher $R^2$ value.

- The improvement is marginal. Moving from the single-parameter M0 model to the three-parameter M3 model only improves the $R^2$ from ~0.93 to ~0.97. This small gain in fitness comes at the cost of tripling the number of fitted parameters.

This pattern of diminishing returns suggests that the additional parameters of the more complex models may not be capturing essential biological mechanisms but rather fitting small amounts of noise in the data.

**Parameter Identifiability Analysis**

To assess confidence in the parameter estimates for our nested models, we performed a Profile Likelihood Analysis (PLA)[2,3]. While goodness-of-fit metrics like AIC can rank models based on their predictive power, they do not indicate whether the parameter values themselves are uniquely determined by the data. PLA provides a rigorous method to quantify this parameter uncertainty and diagnose potential issues of identifiability.

The core principle of PLA is to explore the cost-function landscape for each parameter individually, while allowing all other parameters to re-optimize. For a model with a parameter vector $\theta = (\theta_1, \dots, \theta_p)$, the profile likelihood for a single parameter $\theta_i$ is generated by fixing $\theta_i$ to a specific value and then minimizing the cost function with respect to all other parameters $\theta_{j \neq i}$. This process is repeated across a range of values for $\theta_i$ to trace out its profile. In our case, the cost function is the sum of squared errors ($\chi^2$), so we compute the profile $\chi^2$ function:

$$\chi^2_{\text{profile}}(\theta_i) = \min_{\theta_{j \neq i}} \chi^2(\theta_i, \theta_{j \neq i})$$

A plot of $\chi^2_{\text{profile}}(\theta_i)$ versus $\theta_i$ provides a visual and quantitative measure of the parameter's identifiability:

- A well-identified parameter will exhibit a sharp, parabolic profile with a clearly defined minimum. This indicates that deviations from the optimal parameter value lead to a rapid increase in the cost function, meaning the data provides a strong constraint on the parameter's value.
- A poorly-identified (or non-identifiable) parameter will have a flat or very shallow profile. This indicates that the model's goodness-of-fit is insensitive to changes in the parameter value, meaning the data cannot uniquely determine the parameter.

The 95% confidence interval (CI) for a parameter is determined from this profile. It is defined as the range of parameter values for which the $\chi^2_{\text{profile}}$ is within a certain threshold of its minimum value. This threshold is derived from the $\chi^2$ distribution:

$$\text{CI}_{95\%} = \{\theta_i \mid \chi^2_{\text{profile}}(\theta_i) - \min(\chi^2_{\text{profile}}) \leq \Delta_{0.95}\}$$

where $\Delta_{0.95} \approx 3.84$ for a single parameter (1 degree of freedom).

**Results of PLA**

We performed PLA for the best-fit version of each of the four nested models (M0, M1, M2, and M3), as determined by the lowest AIC score for each model class.

Model M0 contains a single fitted parameter, $k_{syn,shared}$. The profile likelihood plot (SI Figure 3A) shows a clear, parabolic curve with a well-defined minimum. This indicates that the parameter is highly identifiable and well-constrained by the experimental data.

Model M1 adds the volume exponent $n_{shared}$ as a second free parameter. Both $k_{syn,shared}$ and $n_{shared}$ exhibit well-defined, parabolic profiles (SI Figure 3B), although they are shallower than the M0 plot. This demonstrates that even with the added complexity, both parameters remain identifiable, and their optimal values are well-supported by the data.

Model M2 fits separate synthesis rates ($k_{syn,iso}$ and $k_{syn,hypo}$) for the two conditions while fixing the volume exponent to 1 (SI Figure 3C). Here we observe the curves to be even more shallow and parameter like $k_{syn,iso}$ showing poor identifiability in one direction.

Model M3 is the most complex, combining condition-specific synthesis rates with a shared, fitted volume exponent ($n_{shared}$). The PLA for this model reveals a significant issue with parameter identifiability (SI Figure 3d). While the profiles for $k_{syn,iso}$ and $k_{syn,hypo}$ remain reasonably well-defined, the profile for $n_{shared}$ is nearly flat. The $\chi^2$ value changes by less than 1.0 across a wide range of $n_{shared}$ values. This is a classic sign of practical non-identifiability. It suggests a strong correlation between the parameters; the model can achieve a nearly identical goodness-of-fit by compensating for changes in $n_{shared}$ with adjustments to $k_{syn,iso}$ and $k_{syn,hypo}$.

**Parameter Identifiability and Model Selection**

The PLA shows that the parameters for the simplest model M0 is most well-identified with added complexity making the identifiability worse in the most complex models M2 and M3. M3 suffers from practical non-identifiability in its volume exponent parameter, $n_{shared}$. The flatness of its profile likelihood suggests that the model is over-parameterized for the available data. While model selection based on AIC may favor M3 due to its flexibility in fitting the data, the PLA reveals that the specific parameter values of M3 are not reliable. This highlights a trade-off between model complexity and parameter certainty and suggests that the conclusions drawn from the simpler, more identifiable models are more robust.

The marginal benefit of the more complex models is thus challenged by their degree of fit and the PLA done in previous sections. While the parameters of the simpler models (M0, M1, M2) were found to be well-constrained and identifiable, the most complex model (M3) exhibited practical non-identifiability in its volume exponent parameter ($n_{shared}$). This means that the data cannot uniquely determine the value of this parameter in the context of the M3 model, making its value unreliable.

Model M0: a) Provides an excellent fit to the data ($R^2 > 0.93$). b) Captures the core experimental trends. c) Has a single, highly identifiable parameter. Hence, we select the concentration-only model M0. This provides the best support for our core hypothesis that the increased levels of p53RE reporter seen in Hypo-treated cells are due to the size-effect alone, where the smaller nuclear size of Hypo-treated cells causes an increase in the concentration of p53RE.

### Optimization of Data Time Window for Model Fitting

The mathematical models (M0-M3) are formulated to describe the dynamics of p53RE synthesis in response to changes in nuclear volume and p53 concentration. However, the full experimental time course (0-12 hours) contains distinct phases. The early phase (< 3 hours) just after mitosis (after NER) is dominated by rapid, transient changes in nuclear volume. The late phase (> 9 hours) may be influenced by secondary biological processes not included in our model, such as cell cycle progression or complex feedback loops.

Fitting the model across the entire time course could therefore bias the parameter estimates, forcing the model to account for dynamics it was not designed to capture. To obtain the most accurate and reliable parameter values for the core synthesis mechanism, we must identify an optimal time window within the data that best represents the process being modeled.

### Methods for Optimal Window Selection

We employed three main strategies to identify a data window that is both biologically relevant and statistically robust. The start time is guided by a biological anchor, the end time is constrained by a data-driven changepoint analysis, and the final window is selected from a set of candidates using a cross-validation framework.

To isolate the p53RE synthesis dynamics from the initial rapid transients post-mitosis, the analysis window should begin only after the cell's nuclear volume has stabilized. We quantified this by calculating the rate of change of the mean nuclear volume for both Iso and Hypo conditions. The "settling time" was defined as the point at which the rate of volume change fell below a stable threshold. The recommended start time for the analysis window was set to 0.5 hours after the last of the two conditions stabilized, ensuring we exclude the initial adaptation phase.

To find an upper bound for the window, we sought to identify the point at which our model begins to fail in describing the data, suggesting the emergence of confounding late-stage dynamics. We used a method based on the Chow test[4] to detect a structural break in the model's residuals. The model was fit to progressively longer time windows, and for each fit, the residuals (the difference between the model prediction and the data) were calculated. The Chow test was then applied to these residuals to determine if there was a statistically significant difference between the residuals in the first half of the window versus the second half. A significant p-value ($p < 0.05$) indicates a structural break, implying that the model's predictive character changes over the window. The optimal end time was defined as the point just before the first significant structural break occurred.

$$extChowTestStatistic = \frac{(RSS_p - (RSS_1 + RSS_2))/k}{(RSS_1 + RSS_2)/(N_1 + N_2 - 2k)}$$

Where $RSS_p$ is the residual sum of squares for the whole window, $RSS_1$ and $RSS_2$ are the sums for the two halves of the window, $N$ is the number of data points, and $k$ is the number of parameters.

With the start and end times constrained, we performed a cross-validation (CV) analysis to rank a set of high-quality candidate windows. For each candidate, the data was repeatedly split into a training set (e.g., the first 70% of the window) and a validation set (the remaining 30%). The model was fit on the training data, and its ability to predict the unseen validation data was measured by the mean squared error. This process yields a robust CV score for each window, which quantifies its out-of-sample predictive power. A lower CV score indicates a more robust and generalizable model fit.

The final optimal window was chosen by integrating these three analyses. From the list of candidate windows, we selected the one with the lowest CV score that also satisfied the following criteria: 1. Starts at or after the time determined by the Biological Anchor. 2. Ends at or before the time determined by the changepoint analysis. 3. Has a minimum duration of 6 hours to ensure sufficient data for a robust fit.

**Results and Final Window Selection**

We applied this optimization procedure independently to the previously selected model M0. The changepoint analysis identified an acceptable upper time bound of about 9 hours (SI Figure 4A), while the volume dynamics identified the lower bound to be about 3 hours (SI Figure 4B). We also evaluated a set of candidate windows to assess the trade-off between predictive power (CV score, lower is better) and goodness-of-fit (R-squared, higher is better) (SI Figure 4C). The CV scores are shown as bars (left axis), and the R-squared values are overlaid as a line plot (right axis).This plot shows that while several windows achieve a high R-squared value, the CV score (which measures predictive robustness) varies more significantly. The ideal window is one that minimizes the CV score while maximizing the R-squared value. Based on all these analyses, we selected the time window 2-8 hours as optimal for our purposes (SI Figure 4D). The data indicated that the lower bound could be decreased a bit beyond what the volume dynamics calculated which is more conservative


**Supplementary Information Figure Legends**

**SI Figure 1:** Representative model fits for the four nested mathematical models describing p53RE dynamics under iso-osmotic (Iso) and hypo-osmotic (Hypo) conditions. Each panel shows the fit for a specific model: **(A)** Model M0 (concentration-only, 1 parameter), **(B)** Model

M1 (shared volume exponent, 2 parameters), **(C)** Model M2 (condition-specific amplitude, 2 parameters), and **(D)** Model M3 (flexible volume model, 3 parameters). In each panel, the left plot shows Normalized Total p53RE and the right plot shows Normalized Mean p53RE over time. Dotted lines represent experimental data (exp) and solid lines represent the corresponding model fit (model). Red lines correspond to Iso and blue lines to Hypo. Shaded areas represent the standard error of the mean (SEM) for the experimental data and the confidence interval for the model fits. The combined R-squared ($R^2$) and Akaike Information Criterion (AIC) values for each model fit are shown in the inset boxes.

**SI Figure 2:** Sensitivity of model performance metrics to the assumed p53RE degradation rate ($d_{r,shared}$). **(A)** The Akaike Information Criterion (AIC), where lower values are better, against $d_{r,shared}$. **(B)** Combined R-squared ($R^2$), where higher values are better, against $d_{r,shared}$. Each colored line represents one of the four nested models (M0-M3). **(C)** A plot visualizing the trade-off between model goodness-of-fit (R-squared, x-axis) and model complexity penalized by the corrected Akaike Information Criterion (AICc, y-axis). Each point represents a model fit at a specific $d_{r,shared}$, with color and shape indicating the model class (M0-M3). The legend specifies the number of fitted parameters for each model.

**SI Figure 3:** Profile Likelihood Analysis (PLA) for the fitted parameters of each of the four nested models to assess parameter identifiability. For each plot, the solid blue line shows the profile chi-squared ($\chi^2$) objective function as a function of the parameter value. The dashed red line indicates the 95% confidence interval (CI) threshold, with the resulting CI range shaded in pink. The dotted green line marks the maximum likelihood estimate (MLE) for the parameter. **(A)** PLA for the single fitted parameter ($k_{syn,shared}$) of Model M0. **(B)** PLA for the two fitted parameters ($k_{syn,shared}$ and $n_{shared}$) of Model M1. **(C)** PLA for the two fitted parameters ($k_{syn,iso}$ and $k_{syn,hypo}$) of Model M2. **(D)** PLA for the three fitted parameters ($n_{shared}$, $k_{syn,iso}$ and $k_{syn,hypo}$) of Model M3.

**SI Figure 4:** Determination of the bounds for the optimal model fitting window. **(A)** Change-point analysis using the Chow test on the residuals of Model M0 fitted to progressively longer time windows. The p-value dropping below the 0.05 significance level (red dashed line) indicates a structural break, defining the upper bound for the fitting window at approximately 9 hours. **(B)** Normalized nuclear volume dynamics for Iso (red) and Hypo (blue) conditions. The analysis identifies the time point where volume stabilizes post-mitosis, serving as a biological anchor for the start time of the fitting window. The green dashed line indicates the determined recovery start time of 4.0h. **(C)** Comparison of the candidate time windows. Blue bars represent the cross-validation (CV) error (left axis, lower is better), while the orange line plot shows the R-squared value (right axis, higher is better). This analysis identifies the window with the best trade-off between predictive robustness and goodness-of-fit. **(D)** Representative fits of Model M0 to the Iso and Hypo data for two candidate time windows: Candidate 1 ([3.0, 9.0]h) and Candidate 2 ([2.0, 8.0]h).

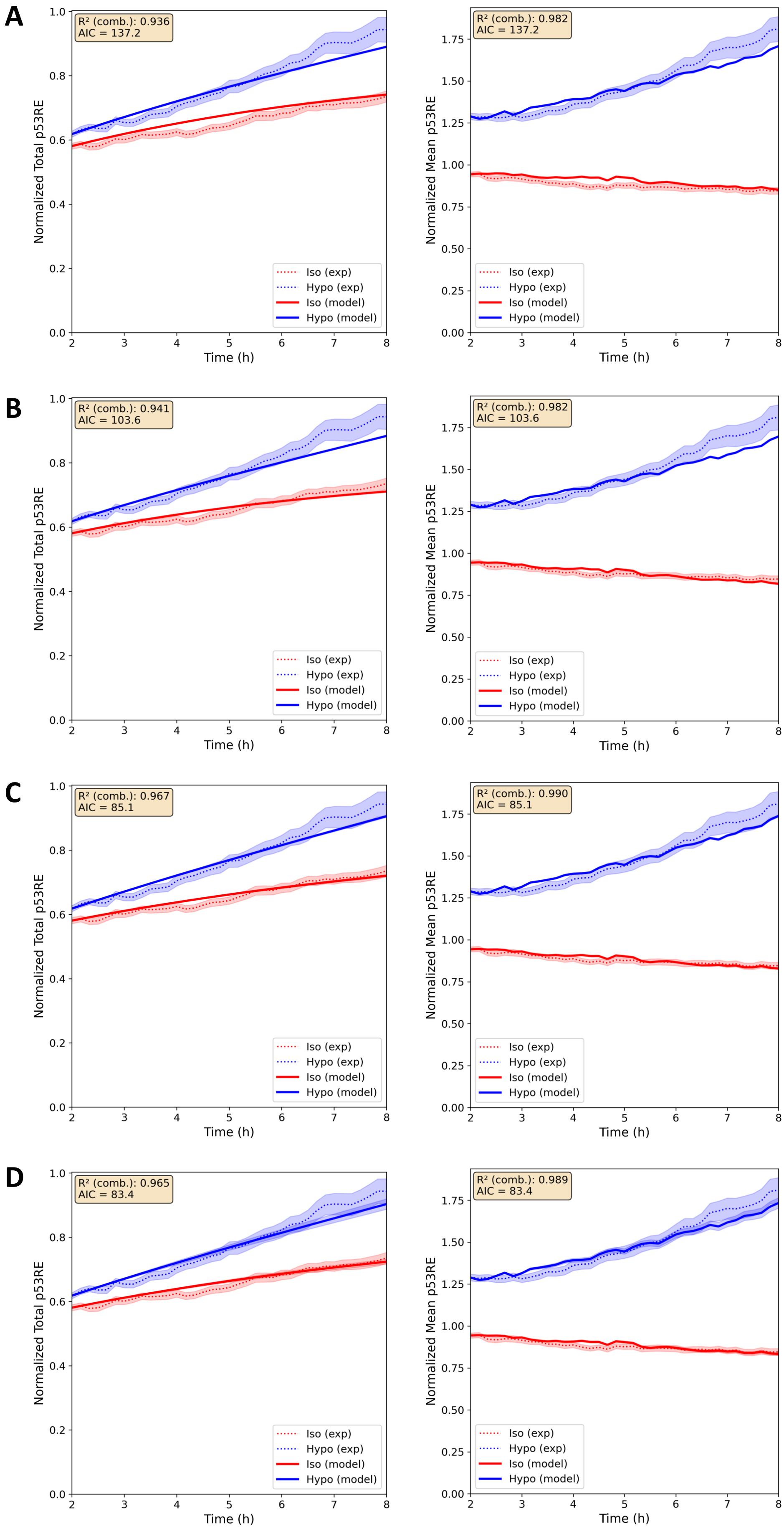
A
R² (comb.): 0.936
AIC = 137.2
Normalized Total p53RE
Time (h)
Iso (exp)
Hypo (exp)
Iso (model)
Hypo (model)
R² (comb.): 0.982
AIC = 137.2
Normalized Mean p53RE
Time (h)
Iso (exp)
Hypo (exp)
Iso (model)
Hypo (model)
B
R² (comb.): 0.941
AIC = 103.6
Normalized Total p53RE
Time (h)
Iso (exp)
Hypo (exp)
Iso (model)
Hypo (model)
R² (comb.): 0.982
AIC = 103.6
Normalized Mean p53RE
Time (h)
Iso (exp)
Hypo (exp)
Iso (model)
Hypo (model)
C
R² (comb.): 0.967
AIC = 85.1
Normalized Total p53RE
Time (h)
Iso (exp)
Hypo (exp)
Iso (model)
Hypo (model)
R² (comb.): 0.990
AIC = 85.1
Normalized Mean p53RE
Time (h)
Iso (exp)
Hypo (exp)
Iso (model)
Hypo (model)
D
R² (comb.): 0.965
AIC = 83.4
Normalized Total p53RE
Time (h)
Iso (exp)
Hypo (exp)
Iso (model)
Hypo (model)
R² (comb.): 0.989
AIC = 83.4
Normalized Mean p53RE
Time (h)
Iso (exp)
Hypo (exp)
Iso (model)
Hypo (model)

SI Figure 1

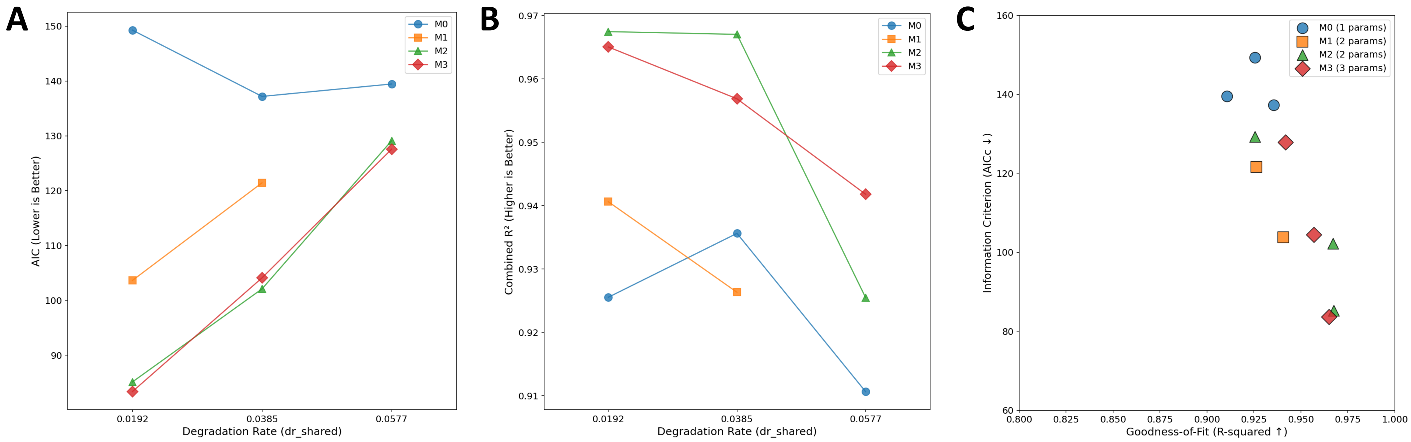
A
AIC (Lower is Better)
Degradation Rate (dr_shared)
M0
M1
M2
M3
B
Combined R² (Higher is Better)
Degradation Rate (dr_shared)
M0
M1
M2
M3
C
Information Criterion (AICc ↓)
Goodness-of-Fit (R-squared ↑)
M0 (1 params)
M1 (2 params)
M2 (2 params)
M3 (3 params)

SI Figure 2

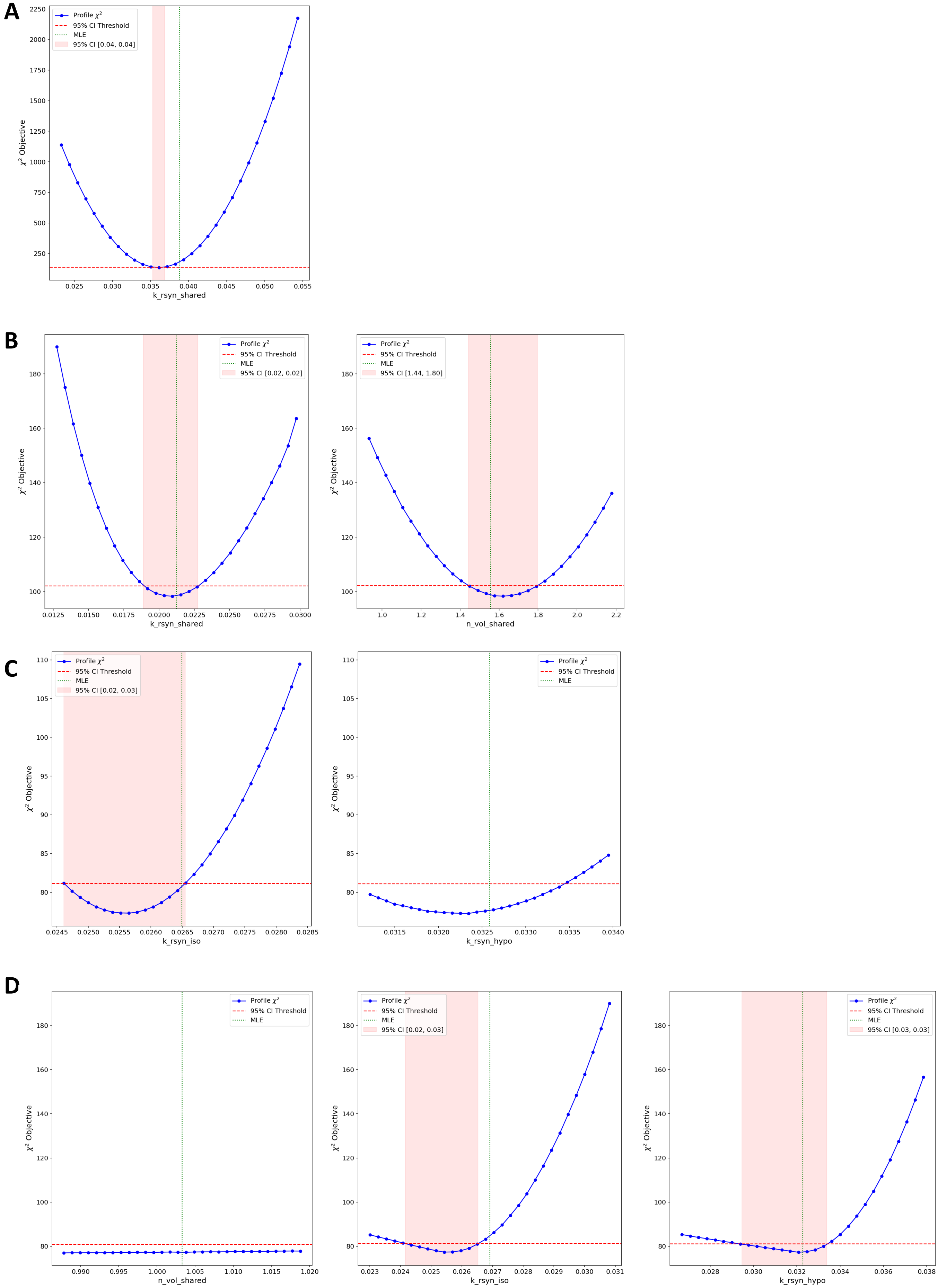
A
Profile $\chi^2$
95% CI Threshold
MLE
95% CI [0.04, 0.04]
$\chi^2$ Objective
k_rsyn_shared
B
Profile $\chi^2$
95% CI Threshold
MLE
95% CI [0.02, 0.02]
$\chi^2$ Objective
k_rsyn_shared
Profile $\chi^2$
95% CI Threshold
MLE
95% CI [1.44, 1.80]
$\chi^2$ Objective
n_vol_shared
C
Profile $\chi^2$
95% CI Threshold
MLE
95% CI [0.02, 0.03]
$\chi^2$ Objective
k_rsyn_iso
Profile $\chi^2$
95% CI Threshold
MLE
$\chi^2$ Objective
k_rsyn_hypo
D
Profile $\chi^2$
95% CI Threshold
MLE
$\chi^2$ Objective
n_vol_shared
Profile $\chi^2$
95% CI Threshold
MLE
95% CI [0.02, 0.03]
$\chi^2$ Objective
k_rsyn_iso
Profile $\chi^2$
95% CI Threshold
MLE
95% CI [0.03, 0.03]
$\chi^2$ Objective
k_rsyn_hypo

SI Figure 3

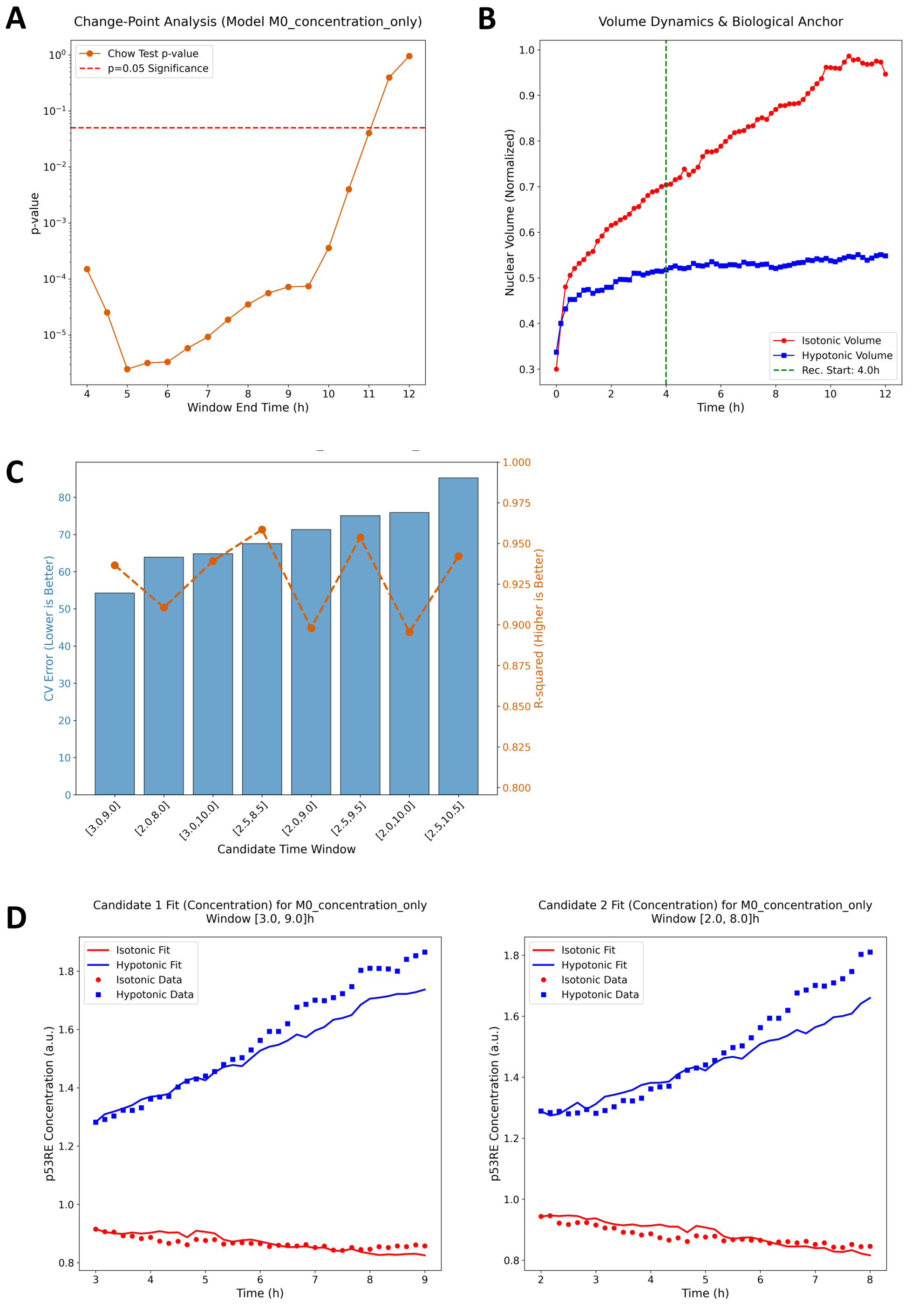

A
Change-Point Analysis (Model M0_concentration_only)
Chow Test p-value
p=0.05 Significance
p-value
Window End Time (h)
B
Volume Dynamics & Biological Anchor
Nuclear Volume (Normalized)
Isotonic Volume
Hypotonic Volume
Rec. Start: 4.0h
Time (h)
C
CV Error (Lower is Better)
R-squared (Higher is Better)
Candidate Time Window
[3.0,9.0]
[2.0,8.0]
[3.0,10.0]
[2.5,8.5]
[2.0,9.0]
[2.5,9.5]
[2.0,10.0]
[2.5,10.5]
D
Candidate 1 Fit (Concentration) for M0_concentration_only
Window [3.0, 9.0]h
Isotonic Fit
Hypotonic Fit
Isotonic Data
Hypotonic Data
p53RE Concentration (a.u.)
Time (h)
Candidate 2 Fit (Concentration) for M0_concentration_only
Window [2.0, 8.0]h
Isotonic Fit
Hypotonic Fit
Isotonic Data
Hypotonic Data
p53RE Concentration (a.u.)
Time (h)


SI Figure 4
